# Optimized Urine Metagenomic Methods Reveal Longitudinal Microbial Community Dynamics and Predictors of Transition from Asymptomatic Colonization to CAUTI

**DOI:** 10.64898/2026.07.06.736792

**Authors:** Namrata Deka, Erin M. Nawrocki, Aimee L. Brauer, Saptarshi Chakraborty, Vaughn S. Cooper, Chelsie E. Armbruster

## Abstract

**Background:** Urinary tract infections (UTIs) rank among the most common infections globally, with many linked to indwelling urinary catheters. Our prior culture-based longitudinal evaluation of long-term catheterized nursing home residents revealed persistent asymptomatic colonization by pathogens and demonstrated that CAUTI onset was not necessarily due to new pathogen acquisition. In this study, we optimized metagenomics methods to examine the ecological structure underlying persistent colonization and the transition to infection.

**Results:** We present a comprehensive longitudinal metagenomic analysis of catheterized urine specimens, revealing colonization dynamics of 69 microbial species across 198 samples from 9 individuals. Descriptive ecological metrics were combined with Bayesian mixed-effects models that accounted for repeated within-participant sampling to identify clusters of co-occurring species, determine the impact of perturbations such as antibiotic exposure and catheter changes on community structure, and identify taxa predictive of infection sign and symptom onset. Longitudinal specimens clustered into three main ecological phenotypes: 1) moderate diversity, unstable communities (3 participants); 2) high diversity, stable communities that resisted disruption even after multiple catheter changes (3 participants); and 3) low diversity, pathogen-dominated communities (3 participants). Catheter changes alone did not significantly disrupt community composition, while antibiotic exposures induced major shifts often followed by re-colonization with the same genera within subsequent weeks. Six clusters of species were identified for which relative abundances correlated across perturbations to the microbial community, including a mutually exclusive Enterobacterales cluster and fastidious-anaerobe group cluster. 24 species were found to correlate with onset of signs and symptoms of infection, 11 of which were missed by standard urine culture.

**Conclusions:** The catheterized urinary tract represents a novel ecosystem that is resilient to disruption by catheter changes but susceptible to antibiotic perturbation. Antibiotic exposure did deplete all species associated with signs and symptoms but also depleted potentially benign microbes. Our findings have direct implications for catheter management protocols and antibiotic stewardship in long-term catheterized patients. Prospective evaluation using this framework in a larger cohort can help translate these ecological insights into clinical decision-making tools.

## INTRODUCTION

Indwelling urinary catheters are ubiquitous in modern healthcare and long-term care settings and are a major driver of urinary tract infections (UTI) [1–3]. Up to 10% of residents in long-term care settings such as nursing homes require permanent catheter use, and these patients typically exhibit continuous bacteriuria with a high risk of infection and severe sequelae [1, 4]. The presence of an indwelling catheter dramatically alters the bladder environment to facilitate bacteriuria, making it the leading cause of healthcare-associated UTIs [1, 4]. However, distinguishing asymptomatic bacteriuria from catheter-associated UTI (CAUTI) can be clinically challenging due to a high prevalence of subjective, nonspecific signs and symptoms of infection [3, 5]. The burden of CAUTI is estimated to be ∼449,334 cases annually in the US alone, leading to prolonged hospitalization, increased morbidity, and healthcare costs exceeding $350 million per year [6, 7]. Urinalysis also provides little diagnostic utility for differentiating CAUTI from bacteriuria due to the inflammation and proteinuria induced by the catheter presence [5]. Together, these problems lead to both insufficient and inappropriate antibiotic use, which highlights the need for improved diagnostic criteria in the long-term catheterized population.

Clinical management of UTIs has historically followed the view that healthy urine is sterile or at least lacking in traditional uropathogens. However, this paradigm has been increasingly challenged with substantial evidence demonstrating the presence of a urinary microbiome [5, 8–10]. Indwelling urinary catheters further create a unique ecological niche that fundamentally alters the urinary tract environment and favors bacteriuria [11]. The catheter surface provides a substrate for biofilm formation, while catheter-associated trauma to the urothelium may disrupt host defense mechanisms. In catheterized individuals, we and others have demonstrated that culturable microorganisms persist as stable, polymicrobial communities resilient to interventions like antibiotics or catheter changes [5, 10]. However, as standard clinical culture methods under-sample fastidious and anaerobic organisms, these data provide only a narrow view of the complete microbiota of the catheterized urinary tract. Furthermore, culture based longitudinal assessments of catheterized urine have revealed that microbes identified during symptomatic CAUTI were often present asymptomatically for several weeks prior to diagnosis, demonstrating that symptom onset is not necessarily driven by the acquisition of a new pathogen but may arise from within a resident microbial reservoir [5, 10].

The clinical significance of polymicrobial communities within the catheterized bladder and their relationship to development of symptomatic disease from asymptomatic colonization remains poorly defined. Longitudinal profiling of catheterized individuals may therefore provide key insights into microbial community dynamics that cross-sectional studies cannot fully capture. Potential mechanisms influencing symptom onset may include the acquisition of new virulent species, the loss of protective commensals, or an increase in the pathogenic potential of resident strains through within-host evolution [5, 12–16]. There is growing evidence for the latter, with studies documenting progressive increases in antibiotic resistance in persistent isolates and the accumulation of resistance genes via horizontal gene transfer within clinical settings [13, 15, 17]. However, most large-scale studies remain culture-based and low-resolution, leaving it unclear whether symptomatic transitions are driven by species-level turnover or by strain-level changes in virulence and resistance.

No prior study has aligned longitudinal whole-genome sequencing of samples from catheterized individuals with infection signs and symptoms. This approach can potentially determine whether symptomatic infection aligns with new species acquisitions, a loss of beneficial microbes, or genomic changes within persistent strains. We therefore conducted a longitudinal metagenomic study designed to evaluate and overcome technical challenges of sequencing complex catheter urine specimens and to characterize all microbes present in the urine. We first optimized methods for microbial DNA extraction and host DNA depletion to maximize resolution of all microbial members in catheter urine specimens. This was followed by metagenomic sequencing of serial urine specimens from chronically catheterized patients to 1) characterize the complete composition of catheterized urine, 2) examine community stability over time, and 3) assess the contribution of specific constituents and community dysbiosis to the development of CAUTI signs and symptoms.

Here we present the first genome-wide characterization of longitudinal urine specimens from 9 catheterized individuals, comprising 198 serial samples collected over 30 weeks. We identified 69 unique species and characterized their temporal dynamics in relation to clinical parameters including catheter changes, antibiotic exposures, and clinical signs and symptoms of CAUTI. Known uropathogens ranged widely in frequency within and among patient samples. Nonetheless patients grouped into distinct community phenotypes, with some individuals maintaining stable commensal-dominated communities even through catheter changes, while others exhibited rapid and frequent shifts in dominant species. Abundance shifts in several species undetectable by standard culture methods were correlated with infection signs and symptoms across multiple CAUTI classification schemes, pointing to candidate predictors of infection onset. The results provide new insights into the ecology of the catheter-associated urinary tract and have important implications for developing microbiome-informed diagnosis, treatment, and prevention for CAUTIs.

## METHODS

### Engineered urine specimen control for DNA extraction and sequencing

To examine the quality of nucleic acid extraction kits and day to day reproducibility, a control urine sample was engineered. The engineered urine control was designed to represent the complexity of a typical clinical urine sample obtained from a patient with a long-term indwelling urinary catheter, including the presence of eukaryotic organisms that could be present in urine. Varying concentrations of four Gram-negative bacteria, two Gram-positive bacteria, two fungi, a protozoan parasite, and a human bladder epithelial cell line were suspended in pooled, filter-sterilized human urine as shown in **Table 1**. 10% human serum was also added to mimic the level of host protein typically present in urine from patients with long-term catheters.

**Table 1:**
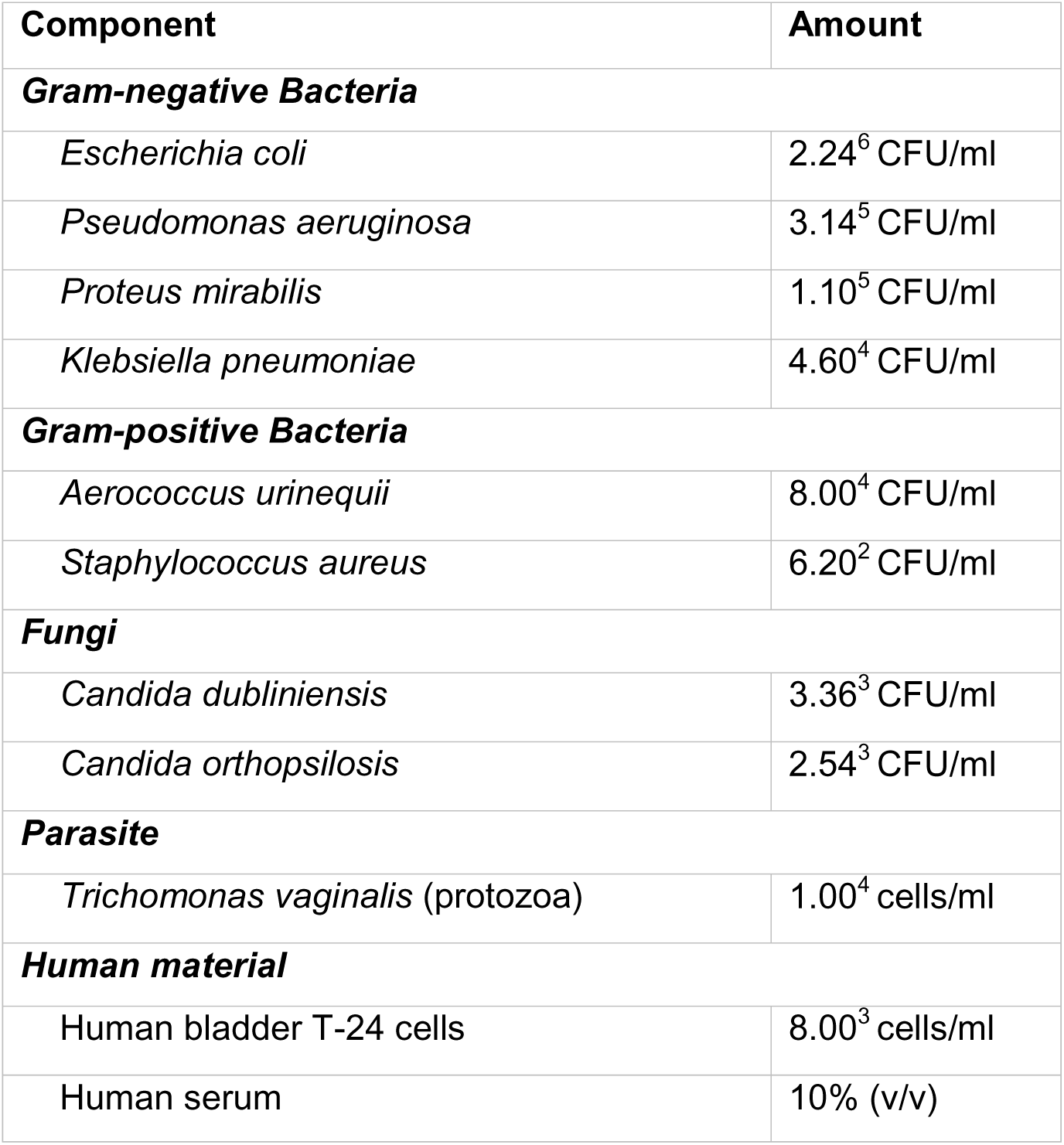
Components in mock urine sample.

### Comparison of methods for microbial DNA extraction from catheter urine specimens

Three microbial DNA extraction kits were selected to evaluate performance for isolating microbial DNA from urine collected from catheterized nursing home participants: 1) Zymo HostZERO Microbial DNA kit (Zymo Research D4310) for select lysing of host material and degradation of extracellular DNA followed by microbial DNA extraction [18], 2) Qiagen QIAamp BiOstic Bacteremia DNA kit (Qiagen 12240-50) for isolation from samples with low bacterial counts, including inhibitor removal [19], and 3) MagMAX Microbiome Ultra Nucleic Acid Isolation Kit (Applied Biosystems by Thermo Fisher Scientific A42354) for efficient nucleic acid extraction from bacteria, fungi, and viruses [20]. All extractions were performed following the manufacturer’s instructions, with specific modifications noted below to improve DNA yield. An initial hypotonic lysis step was explored in combination with the QIAmp and MagMAX extraction methods to remove host material prior to complete lysis of microbes and was adapted from [21].

#### Hypotonic Lysis

250μL of urine sample was suspended in 7mL dH_2_O and incubated at room temperature on nutating mixer with gentle agitation for one hour. 800μL of Benzonase buffer (200 mM Tris-HCl, 10 mM MgCl_2_) and 250U Benzonase (Sigma E-1014) was added to sample and incubated at 37°C for 2 hours with gentle agitation. The sample was spun at 8000g for 10 minutes, the pellet was washed once in 1X PBS and then resuspended in 400 μL TE buffer. The Benzonase reaction was quenched by adding 88 μL of 0.5M EDTA and 880 μL of 1.5M NaCl and sample could then proceed with nucleic acid extraction kit.

#### Zymo HostZERO Microbial DNA kit

Two replicate urine aliquots were processed and combined where indicated during the microbial DNA isolate step. For host DNA removal, 250 µL of each urine sample was added to 1 mL Host DNA Depletion Solution and incubated at room temperature on a nutating mixer with gentle agitation for 15 minutes. Samples were centrifuged at 10,000g for 5 minutes, supernatants carefully removed, and pellets resuspended in 100 µL Microbial Selection Buffer. 1 µL of Microbial Selection Enzyme was then added to each suspension, briefly vortexed to mix, and incubated at 37°C for 30 minutes. The samples then underwent the manufacturer’s recommended Proteinase K treatment by adding 20 µL Proteinase K, vortexing for 10 seconds, and incubating at 55°C for 10 minutes. Following incubation, 100 µL DNA/RNA Shield was added, vortexed for 10 seconds, and incubated at room temperature for 10 minutes.

For microbial DNA isolation, the entire sample from the host DNA depletion step was added to a ZR BashingBead Lysis Tube. 750 µL of ZymoBIOMICS Lysis Solution was added to the tube, tubes were secured in a vortex tube holder, vortexed at maximum speed for 10 minutes, and centrifuged at 10,000g for one minute. The supernatant was transferred to a collection tube and 1,200 µL of ZymoBIOMICS DNA Binding Buffer was added and mixed well. At this step, the two replicate tubes from the same urine sample were combined and the supernatant/DNA binding buffer mix was added to a Zymo-Spin IC-Z Column in a collection tube and centrifuged at 10,000g for 30 seconds and flow-through was discarded. Columns were reloaded and spun again until all supernatant/DNA binding buffer was added. 400 µL of ZymoBIOMICS DNA Wash Buffer 1 was added to column and centrifuged at 10,000g for 30 seconds. 700 µL of ZymoBIOMICS DNA Wash Buffer 2 was added to the column and centrifuged at 10,000g for 30 seconds. Lastly, 200 µL of ZymoBIOMICS DNA Wash Buffer 2 was added to the column and centrifuged at 10,000g for one minute for complete removal of wash buffer. The column was then transferred to a clean microcentifuge tube and 20 µL of ZymoBIOMICS DNase/RNase-Free water prewarmed to 55°C was added directly to column and allowed to sit for 10 minutes. The column was centrifuged at maximum speed for one minute to elute DNA. The eluted DNA was returned to column and allowed to sit for an additional 10 minutes before centrifuging again to ensure maximum retrieval of DNA.

#### Qiagen QIAamp BiOstic Bacteremia DNA kit

The starting volume was increased to 500 µL of each urine specimen to increase DNA yield. Additionally, only 20 µL of Solution EB was used to elute DNA, and the elution was run over the column twice to increase DNA yield.

#### MagMAX Microbiome Ultra Nucleic Acid Isolation Kit

The starting volume was increased to 500 µL of each urine specimen to increase DNA yield. Samples subjected to hypotonic lysis were pelleted and resuspended in 800 µL Lysis Buffer prior to start. Elution Solution provided with kit was prewarmed to 55°C and only 20 µL was used to elute DNA off beads to increase yield.

### Microbial DNA extraction from long-term catheterized urine specimens

Serial urine samples were previously collected at sequential weekly clinical visits for 19 participants [5]. A prospective observational cohort study of asymptomatic catheter-associated bacteriuria was conducted across two Buffalo-area nursing homes between July 2019 and March 2020. Eligible residents had an indwelling urinary catheter (Foley or suprapubic) for at least 12 months, were at least 21 years of age, and could provide informed consent; residents receiving end-of-life care were excluded. Participants were followed weekly for up to 7 months, with each visit comprising chart review, clinical symptom assessment, and urine specimen collection by licensed nurses. Participants were withdrawn upon request, catheter removal without replacement, facility transfer, or death. Baseline chart reviews captured participant demographics, comorbidities, functional status, catheterization history, and prior antimicrobial use. Weekly chart reviews recorded suspected infections, hospitalizations, urine culture and urinalysis results, and antimicrobial prescriptions. Signs and symptoms of CAUTI assessed at each visit included fever, suprapubic or costovertebral pain or tenderness, hypotension, chills or rigors, and acute mental status change. Medical records from any interim hospitalizations were incorporated into the dataset. All study data were managed using REDCap [22–24] hosted through the University at Buffalo Clinical and Translational Science Institute.

In this study, we sequenced all the longitudinal urine samples collected for 9 participants who completed at least 19 study visits (range 19-30 weeks). DNA extraction using the Zymo HostZERO Microbial DNA kit (Zymo Research D4310) was found to provide an abundance profile that most closely resembled the expected abundance of the engineered urine specimen. This kit was therefore selected for extraction of DNA from all longitudinal urine specimens. 1ml of urine was supplemented with Zymo urine conditioning buffer and stored at -80 for DNA extraction. For each DNA extraction, an engineered urine specimen aliquot and water control were also subjected to the extraction process and sequenced.

### DNA sequencing of clinical samples

Extracted DNA was subjected to shotgun metagenomics. Sequencing libraries were constructed using the purePlex DNA Library Prep Kit (seqWell, Beverly, MA). Libraries were pooled and quantified with Qubit High Sensitivity dsDNA reagents, and fragment sizes were assessed on Agilent TapeStation D5000 ScreenTape. Pooled libraries were denatured with 0.2 N NaOH, and the average fragment size was used to dilute each pool to a loading concentration of 1.5-1.6 pM in HT1 buffer. The pools were spiked with 1% PhiX and sequenced on an Illumina NextSeq 500 (2 x 150 bp reads). All raw reads are publicly available in NCBI BioProject: PRJNA1466380.

### Bioinformatic analysis

Reads were demultiplexed with Illumina’s bcl2fastq software v2.17.1. FASTQ files were processed with bioBakery’s “wmgx” workflow for whole-metagenome shotgun sequencing using default settings [25]. Briefly, adapters and repetitive sequences were trimmed and reads that aligned to the human genome (hg37dec_v0.1) were removed with KneadData v0.7.6. During pre-filtering, 204 samples passed the minimum read threshold of >1,000 raw reads. Mean raw reads per sample is ∼2.88 million (range: 17,400 – 13.08 million) paired reads. After trimmomatic-based quality filtering, mean reads retained ∼2.39 million per sample. After human genome decontamination using hg37dec_v0.1, mean reads available for microbial profiling were ∼658,800 per sample which removed an average of 72.6% of total reads that were of human origin. The remaining ∼27.4% of sequenced reads were used for taxonomic classification. The remaining nonhuman reads were assigned to microbial taxa in the CHOCOPhlAn database (mpa_v31_CHOCOPhlAn_201901) using MetaPhlAn v3.0.7 [26, 27]. A total of 69 unique bacterial and fungal species were identified across all samples and classified into three ecological categories including 1) uropathogens (n=18): established causative agents of urinary tract infections, 2) oral species (n=16): species whose primary ecological niche is the oral cavity, and 3) non-oral commensal species (urogenital, vaginal tract, skin, gut, environmental, n=32): species that colonize body sites and are not typically associated with infections (**Table S1**) [28, 29]. Species classification was performed using established microbiological and clinical literature, the Human Oral Microbiome Database (HOMD;[28]), BacDive [29], and clinical microbiology references [30–32]. We classified the species based on primary habitat only, because some species (e.g., *Streptococcus anginosus* group) can colonize multiple body sites, and their classification may require individualized consideration based on strain-level characteristics. The ecological classification is summarized in **Table S2**.

Taxonomic profiles were cross-validated for participant 201 by comparing MetaPhlAn4 against Kraken2 v2.1.2 (RefSeq/PlusPF database), applying a stringent >0.5% relative abundance threshold to Kraken2 output to focus on biologically meaningful taxa and mitigate k-mer artifacts [33, 34]. This threshold was validated against an engineered positive-control community (which yielded 89 false-positive species at lower thresholds) and matched the expected species count, with 492 of 543 total species across the dataset never exceeding 0.1% relative abundance and thus classified as artifacts.

### Statistical Analyses

Descriptive non-parametric tests were used for participant-level cross-sectional comparisons, whereas Bayesian mixed-effects models were used for longitudinal exposure and outcome inference. Kruskal-Wallis tests were used to compare participant-level alpha-diversity summaries across participants, and Spearman rank correlation was used to assess the association between median richness and median diversity metrics. Linear regression with Pearson correlation was retained for visualization only, with the regression line shown to illustrate trend direction rather than for inferential testing. Statistical significance for these descriptive analyses was assessed at p = 0.05.

To formally evaluate associations between clinical exposures (antibiotic use and catheter change), microbial community features, and clinical outcomes while accounting for repeated longitudinal sampling within participants and the small cohort size (n = 9), four Bayesian mixed-effects analyses were performed. Models were fit using rstanarm/Stan with weakly informative priors for parts 1-3 and a student-t ridge-like shrinkage prior for part 4. Statistical significance in the Bayesian analyses was assessed using the local false sign rate (lfsr; defined for a coefficient as the minimum of the posterior probabilities of the coefficient being positive and negative), with lfsr < 0.05 considered significant. Posterior summaries are reported as median estimates with 90% (in conjunction with lfsr < 0.05) and 95% equal-tailed credible intervals.

#### Part 1

Richness, Shannon diversity, Simpson diversity, log-transformed Aitchison distance, and logit-transformed Bray-Curtis dissimilarity between consecutive visits were each modeled as outcomes in Bayesian linear mixed-effects models with participant-specific random effects and fixed effects for antibiotic use, catheter change, and their interaction. Participant-specific random effects were included in all longitudinal models. As a sensitivity analysis, PERMANOVA was also performed on Aitchison and Bray-Curtis distance matrices with permutations stratified by participant.

#### Part 2

Centered log-ratio (CLR)-transformed abundances for 69 taxa were modeled jointly in a multilevel linear mixed-effects framework with participant-level and species-level random effects and fixed effects for antibiotic use, catheter change, and their interaction.

#### Part 3

Adjusted species-species associations were estimated from posterior residual CLR values after removing participant and clinical exposure effects, and posterior Pearson and Spearman correlation matrices were summarized after setting coefficients with lfsr > 0.05 to zero.

#### Part 4

Logistic mixed-effects models related four clinical outcome definitions to standardized CLR species abundances and participant-level random intercepts using a Student-t ridge-like shrinkage prior to stabilize estimation in the high-dimensional species model. The clinical outcome definitions were: 1) potential CAUTI based on Infectious Disease Society of America (IDSA) criteria [30], 2) potential CAUTI based on IDSA criteria but only considering new-onset signs and symptoms [30], 3) potential CAUTI based on National Healthcare Safety Network (NHSN) criteria [35], and 4) potential CAUTI based on NHSN criteria but allowing for inclusion of polymicrobial urine cultures [35].

Bray-Curtis dissimilarity was used in three distinct ways in this study: descriptively to flag major ecological shifts (BC > 0.5), as a continuous logit-transformed outcome in Bayesian mixed-effects models, and as input to PERMANOVA sensitivity analyses. All descriptive analyses were conducted in Python v3.9.6 using NumPy, Pandas, SciPy, and Matplotlib, and Bayesian models were fit in R using rstanarm for Stan-based Markov chain Monte Carlo sampling (4 chains; 1,000 warmup and 1,000 post-warmup iterations per chain).

The Bayesian models in Parts 1-3 used default weakly informative priors as implemented in rstanarm, whereas Analysis 4 used a Student-t-based ridge prior because of the high-dimensionality of predictors (69 correlated species predictors were modeled against four clinical outcomes in a cohort of 9 participants). All scripts used in this study can be found in **Additional File 1-scripts.R.**

## RESULTS

### Optimization of DNA extraction methods

To establish a reproducible protocol for metagenomic sequencing of complex catheter urine specimens, we first evaluated DNA extraction and host depletion methods using an engineered mock urine community. This community contained defined quantities of Gram-positive and Gram-negative bacteria, fungi (*Candida* spp.), a parasite (*Trichomonas vaginalis*), and human cellular material to mimic the composition of clinical catheter urine samples (**Table 1**).

Five different DNA extraction methods were compared to identify the best extraction kit and the utility of depleting host DNA prior to extraction, as described in detail in the methods. Hypotonic lysis-based methods inconsistently depleted host DNA and introduced taxonomic bias, notably over-representing *Staphylococcus* (**Fig. 1, Table S3**). The QIAamp kit skewed community representation toward *Staphylococcus*, while the MagMAX kit over-represented *Pseudomonas* and performed poorly for fungal detection. Neither of these methods performed well to detect the parasite (**Fig. 1, Table S3**).

**Fig. 1.**
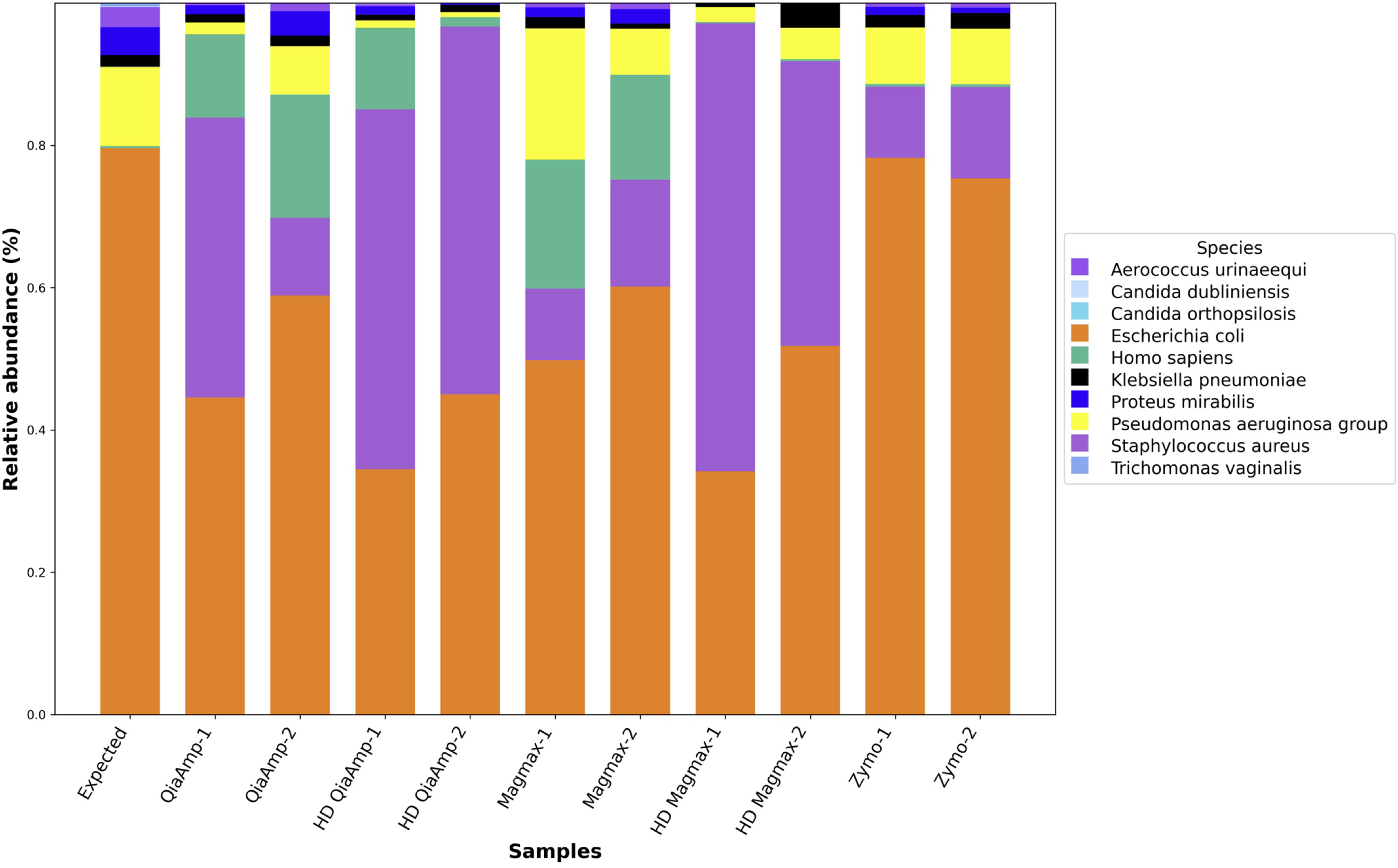
Comparing different DNA extraction kits for reproducible results. The Zymo HostZERO kit best represented the expected frequencies of the engineered microbial community in urine.

In contrast, the Zymo HostZERO kit yielded the highest DNA concentrations, achieved optimal depletion of human DNA, and consistently recapitulated the known microbial composition of the mock community across replicates, despite very limited representation of the parasite (**Fig. 1**). Hence, the Zymo HostZERO method was selected for extracted DNA from the complete longitudinal series of urine specimens from 9 study participants. Water-only and mock community controls were also extracted and processed in parallel, confirming the absence of significant contamination and the continued robustness of the extraction and sequencing pipeline.

### Catheterized urine samples are largely dominated by uropathogens

Across 198 urine samples from 9 participants with long-term catheters, 51.6% of taxa could be categorized as uropathogens, followed by 30.7% as commensal species, and 16.2% as species primarily found in the oral cavity (**Fig. S1a**). When examining community composition dynamics across all isolates, 7/9 participants (78%) had uropathogen dominated profiles (Participants 101, 106, 201, 203, 204, 206, and 207) while 2/9 had commensal-dominated profiles (Participants 102 and 104) (**Fig. S1b**). Oral species were detected in samples from 6/9 participants, while participants 101, 106, and 203 had no detectable oral species with follow up. Participant 104 had the highest overall abundance of oral species across longitudinal samples (44.5%) followed by 206 (41.1%), with a moderate range of 0.1-29.1% abundance observed in rest of the participants (**Fig. S1c**).

### Microbial community structure in catheterized urine is participant-specific

Microbial community structure was specific to each patient in our study (**Fig. S2, Table S4**). The observed number of microbial taxa varied across participants where median richness ranged from 2 to 27 taxa (**Fig. S2a**). Participant 104 exhibited the highest richness, with values consistently clustering around 25 species (range 10-34). In contrast, Participant 203 showed the lowest richness, averaging 3 species (range 1-5). The remaining participants fell into an intermediate band of approximately 8 species, with modest within-participant variability (**Table S4**).

Longitudinal specimens clustered into three main ecological phenotypes: 1) moderate diversity, unstable communities (participants 102, 106, and 204); 2) high diversity, stable communities that resisted disruption even after multiple catheter changes (participants 104, 206, and 207); and 3) low diversity, pathogen-dominated communities (participants 101, 203, 201). A detailed narrative overview of colonization trends and key taxa is provided in **Supplementary Note A** and discussed in brief below.

Three participants (102,106, 204) showcased patterns of transient community disruption. Participant 102, a 61-year-old female with a Foley catheter, exhibited a stable, moderately diverse microbial community was largely maintained over 30 weeks and resilient to repeated catheter changes (Kruskal-Wallis, p = 0.2660), except for a transient episodic bloom of *Enterococcus faecalis* followed by *Proteus mirabilis* between weeks 10–14. The community then reset to its original, *Aerococcus urinae*-dominated structure, illustrating ecological resilience (**Fig. 2a–b**). Participant 106, a 61-year-old male with a suprapubic catheter, exhibited a community structure that was initially resilient to catheter changes but ultimately failed to recover, resulting in a bloom of *Proteus mirabilis* following the week 17 catheter change and a 45% collapse in α-diversity (H: 1.47 → 0.81) by week 19. Similarly, when *P. mirabilis* abundance receded, the community did not return to its original composition but transitioned to a new *Aerococcus urinae*-dominated, low-diversity state (**Fig. 2c–d**). Participant 204, a 51-year-old male with a suprapubic catheter, demonstrated two temporally distinct community states separated by a catheter change at week 13. Prior to week 13, the community was dominated by *Streptococcus anginosus* group with a polymicrobial background of common catheter-associated uropathogens (*Morganella morganii, Streptococcus aureus*) The catheter changes shifted species richness from 10 to 16 within one week. *S. anginosus* group was displaced by *Klebsiella pneumoniae*, a nosocomial uropathogen that persisted as the dominant taxon through end of follow-up (**Fig. 2e–f**).

**Fig. 2.**
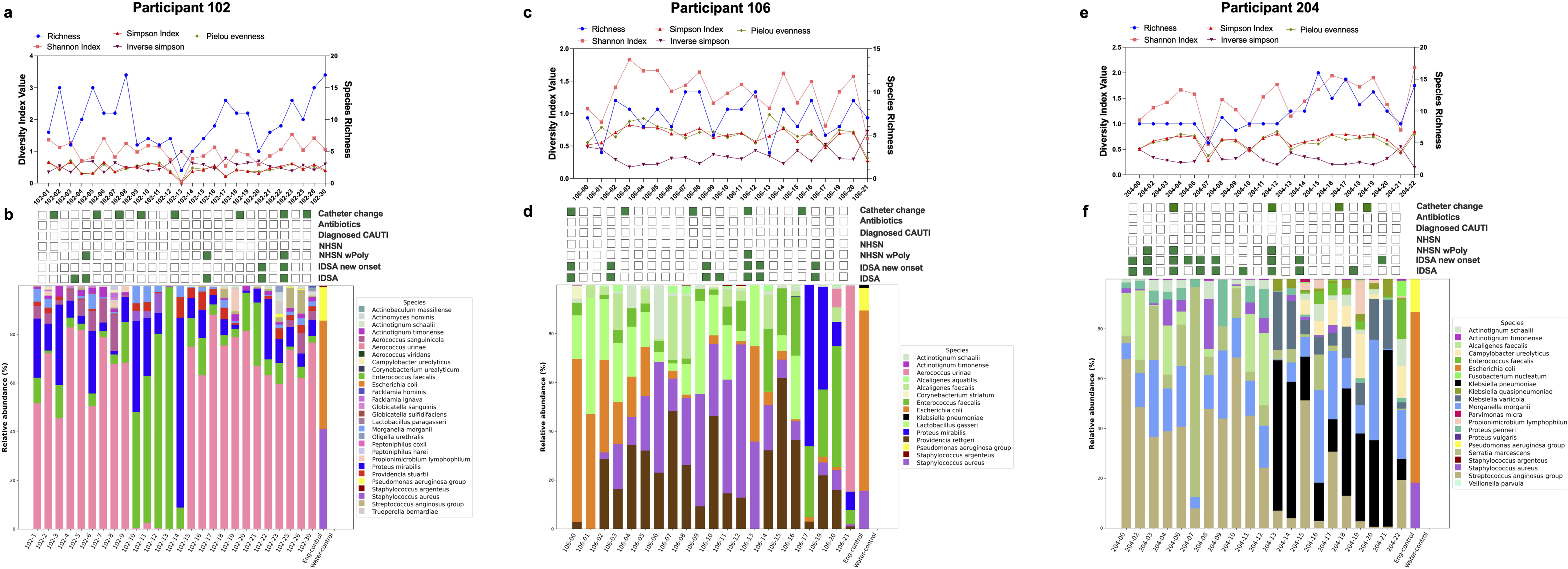
Taxonomic restructuring of the urinary microbiome following catheter change. **2a**. Alpha diversity metrices in Participant 102. **2b**. Stacked bar plot showing the relative abundance of major bacterial taxa across weeks in participant 102. **2c.** Alpha diversity metrices in Participant 106. **2d.** Relative abundance in Participant 106. **2e.** Alpha diversity metrices in Participant 204. **2f.** Relative abundance in Participant 204.

In contrast, participants 104, 206, and 207 maintained broadly stable community structures throughout follow-up, yet each exhibited a distinct ecological signature in taxonomic composition and response to perturbation. Participant 104, a 91-year-old male with a suprapubic catheter, showcased a stable and diverse microbial community over 26 weeks, with α-diversity indices remaining consistently elevated despite 11 catheter changes (S = 18–38; H = 1.7–2.7). The community was dominated by an oral anaerobe consortium including *Propionimicrobium lymphophilum*, *Parvimonas micra*, *Peptococcus niger*, and *S. anginosus* group co-occurring at near-equal relative abundances throughout follow-up. 15 species detected in this participant were absent from the participants with dynamic communities (102, 106, 204). 11/15 were oral anaerobes, while 7 canonical uropathogens found in above were entirely absent from participant 104 structuring an oral anaerobe-dominated community architecture (**Fig. 3a–b**; **Table 2–3, Table S2**). Participant 206, a 44-year-old female with a suprapubic catheter maintained a stable, species-rich community throughout 22 weeks of follow-up. Diversity indices remaining consistently elevated across two catheter changes (H’ = 1.25–2.17; D = 0.51–0.87). The community was characterized by co-dominance of metabolically synergistic organisms including *Veillonella parvula*, a lactate-fermenting oral commensal with recognized uropathogen *M. morganii*, *P. stuartii*, *Pseudomonas aeruginosa,* and low persistence of *P. mirabilis* (**Fig. 3c–d).** Participant 207, a 57-year-old female with a Foley catheter, exhibited similar stable species richness (S = 6–10) over 17 weeks, with notable fluctuations in community evenness, showing a structural reorganization within a persistent core polymicrobial background rather than species turnover. The community was resilient to catheter changes at weeks 3 and 14 but underwent a sustained post-week-14 decline in evenness without species loss (J’= 0.73→0.45) with dominance by a subset of the resident taxa. The core community included *S. aureus*, *E. faecalis*, and *Actinomyces hominis* with oral commensal *Fusobacterium nucleatum* and *Lactobacillus rhamnosus*. This was the only female participant in the cohort with a *Lactobacillus* strain (**Fig. 3e–f**). The list of all the dominant species at each timepoint for the unstable and the stable communities are listed in **Table S4**.

**Fig. 3.**
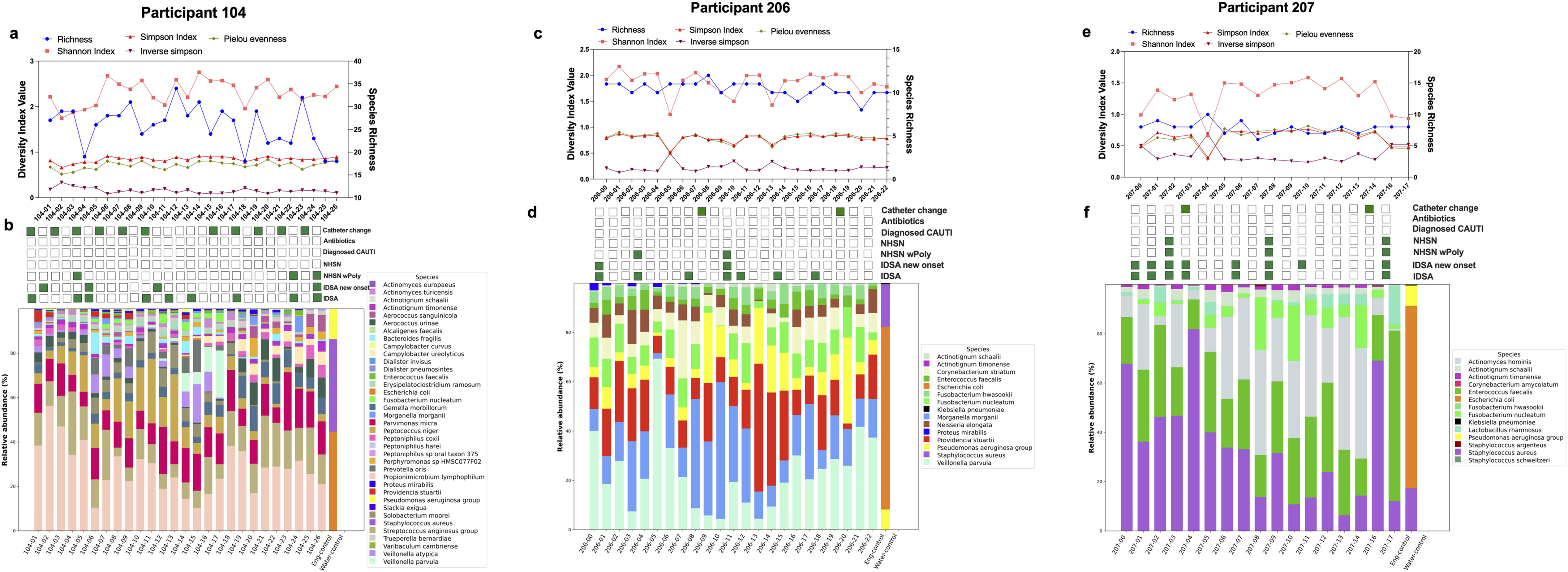
Stable taxonomic structure of the urinary microbiome following catheter change. 3a. Alpha diversity metrices in Participant 104. **3b.** Stacked bar plot showing the relative abundance of major bacterial taxa across weeks in participant 104. **3c.** Alpha diversity metrices in Participant 206. **3d.** Relative abundance in Participant 206. **3e.** Alpha diversity metrices in Participant 207. **3f.** Relative abundance in Participant 207.

**Table 2.**
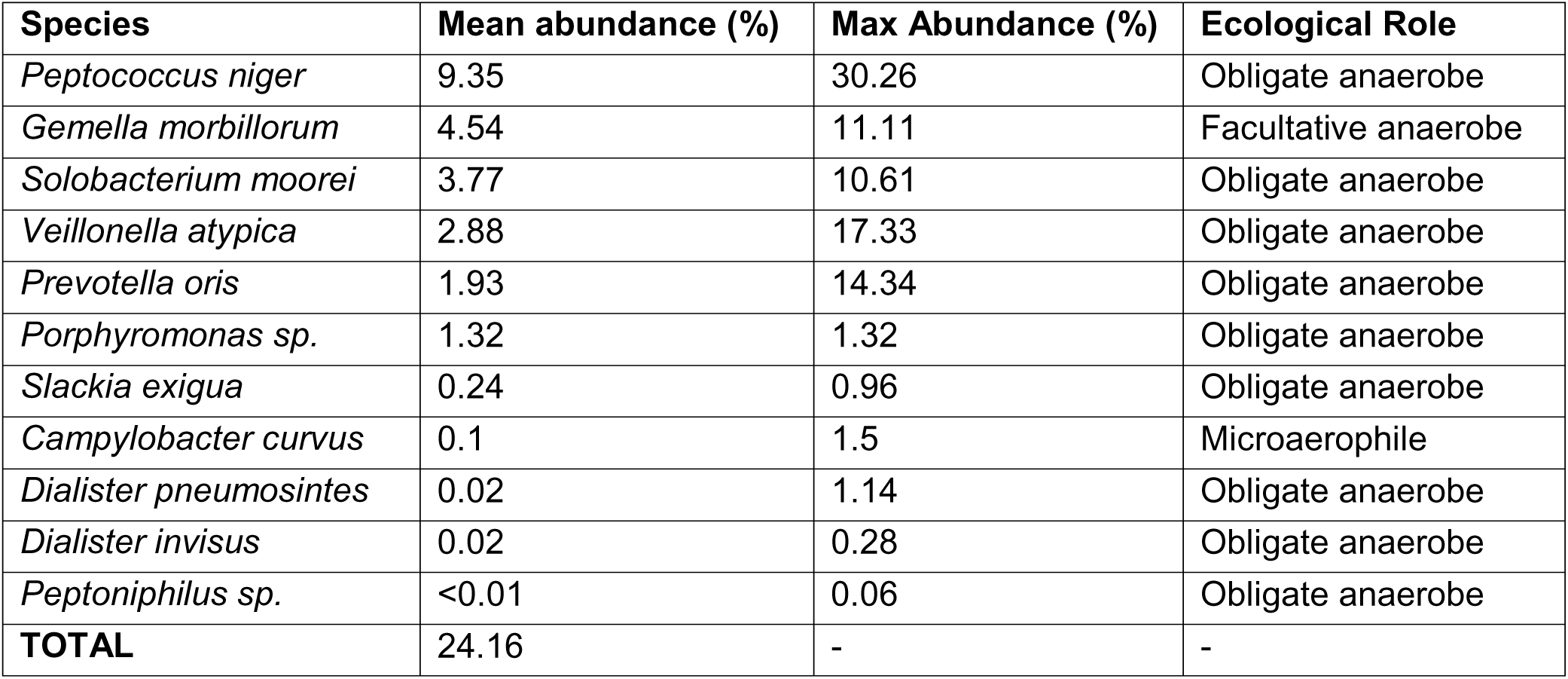
Oral anaerobes unique to the stable urinary microbiome (Participant 104)

**Table 3.**
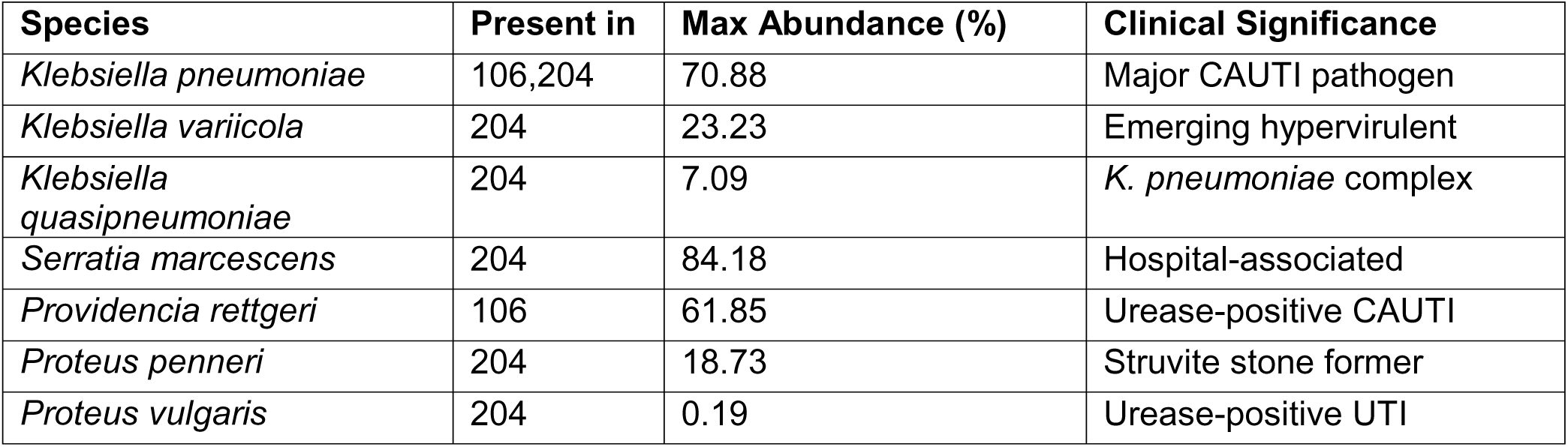
Uropathogen species present in unstable communities (102, 106, 204) but absent from stable community 104.

The remaining three participants exhibited uropathogen-dominated communities with drastic, antibiotic-mediated perturbations to community structure. Participant 101, a 77-year-old male with a suprapubic catheter, exhibited a community that was unperturbed by six catheter changes (Kruskal-Wallis, p = 0.7342), but prophylactic Bactrim (trimethoprim-sulfamethoxazole) administered between weeks 8–9 drove a near-total community collapse, with H’ falling from 1.18 to 0 and richness dropping from 8 species to a single taxon within one week. Antibiotic treatment eliminated TMP-SMX–sensitive bacteria but selected for resistant *P. aeruginosa* (**Table 4**). Diversity recovered to pre-antibiotic levels by week 18 but the community re-established into a *Propionimicrobium lymphophilum* and *P. stuartii* dominated state which was compositionally different from the original *Aerococcus urinae* baseline (**Fig. 4a–b**). Similarly, participant 203, a 68-year-old male with a suprapubic catheter, harbored a persistently low-diversity community (S ≈ 2; H’ ≈ 0.05) for which repeated oral cephalexin administration eliminated *P. mirabilis,* while intrinsically cephalosporin-resistant *P. aeruginosa* remained dominant, similar to the trend observed in participant 101. Each post-antibiotic window was followed by opportunistic blooms including three *Klebsiella* species following final treatment (**Fig. 4c–e**).

**Fig. 4.**
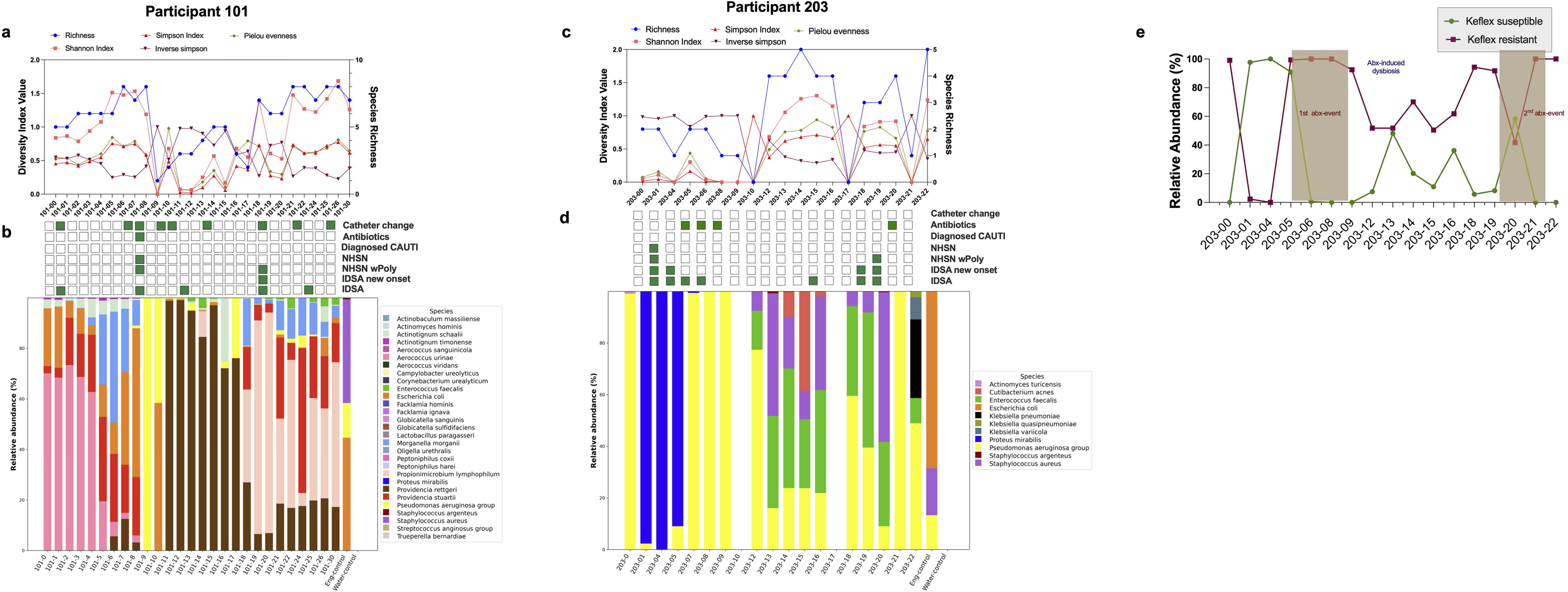
Taxonomic restructuring of the urinary microbiome following catheter change and antibiotic administration. 4a. Alpha diversity metrices in Participant 101. **4b.** Stacked bar plot showing the relative abundance of major bacterial taxa across weeks in participant 101. **4c.** Alpha diversity metrices in Participant 203**. 4d.** Relative abundance in Participant 203**. 4e.** Relative abundance (%) of Keflex (cephalosporin) sensitive and resistant organisms.

**Table 4:**
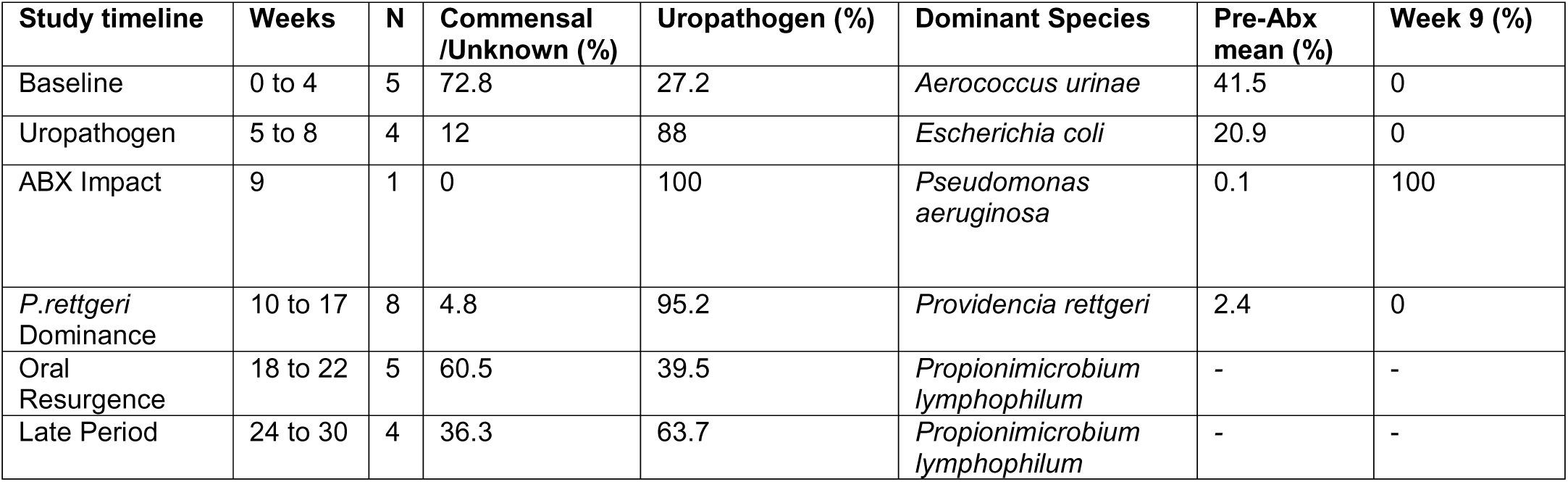
Variation in bacterial relative abundances in response to commonly used antibiotics (trimethoprim and sulfonamide) in UTI.

Participant 201, a 59-year-old male with a suprapubic catheter was the only participant to experience clinician-diagnosed CAUTI in our cohort. Across 7 catheter changes and 3 antibiotic events, both infection episodes arose not from acquisition of a new microbe but from resident taxa present asymptomatically for weeks. The first ciprofloxacin administration induced a transient collapse (S = 7 → 3; H’ = 1.49 → 0.56) from which the core *P. stuartii*, *Peptoniphilus harei*, *S. anginosus* consortium rebounded by week 7 **(Fig. S3)**. The second, severe episode (pyelonephritis with sepsis one day after the week 17) was preceded by co-dominance of *P. stuartii* (45.1%) and *M. morganii* (10.7%) **(Fig. S3)**. Antibiotic treatment drove a 21-fold collapse of this pathogen pair to 2.6% while *S. anginosus* rose to 54.6% as the dominant successor, with all diversity metrics reaching zero by week 19 (**Fig. 5a–b**). Because antibiotic exposure in participants 201 and 101 was specifically UTI- associated compared to other participants, their composition is informative for the ecological consequences of treatment on catheter-associated microbiota.

**Fig. 5.**
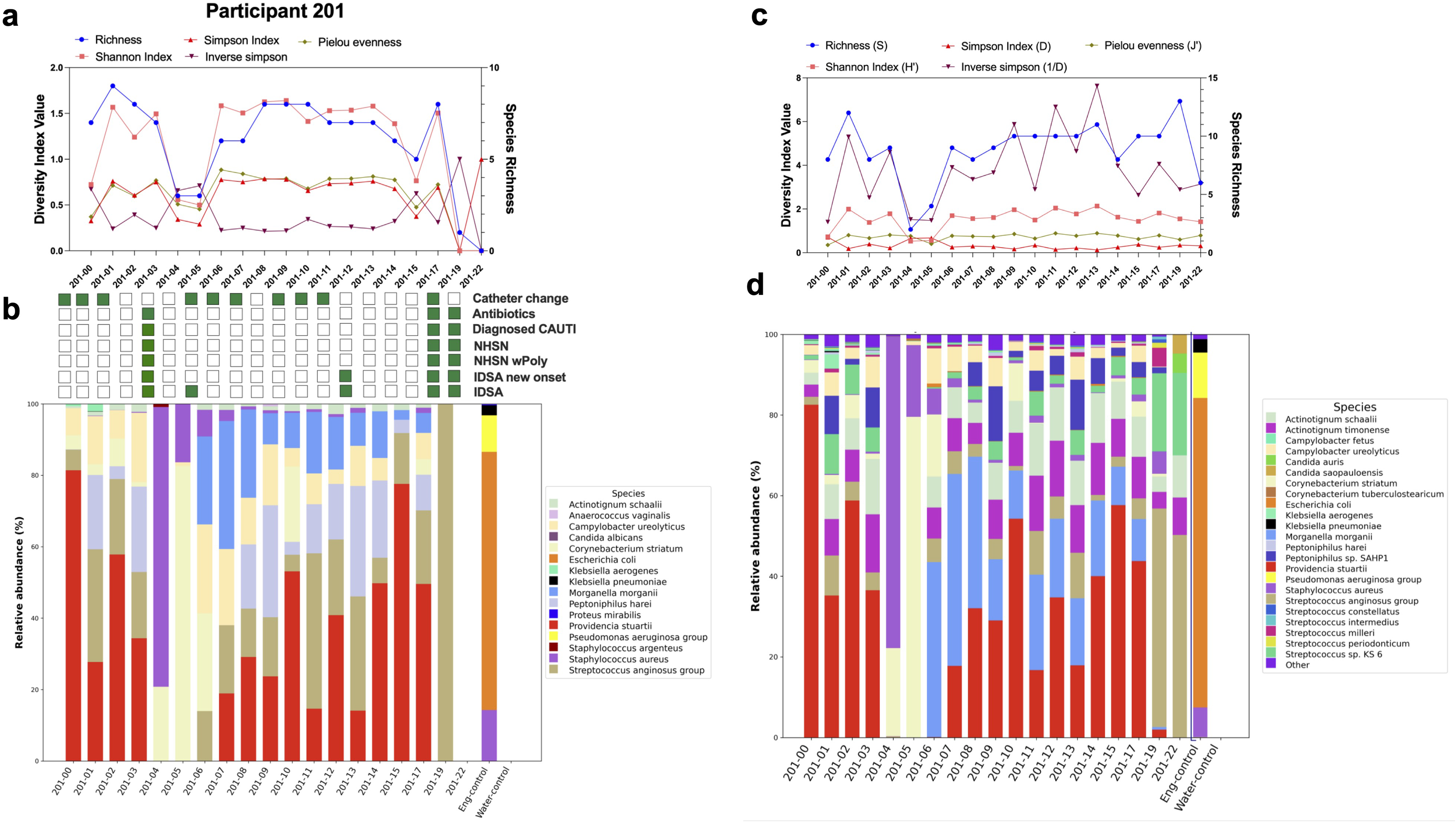
Taxonomic restructuring of the urinary microbiome following catheter change and antibiotic administration. 5a. Alpha diversity metrices in Participant 201**. 5b.** Stacked bar plot showing the relative abundance of major bacterial taxa across weeks in participant 201**. 5c.** Kraken 2 alpha diversity metrices in Participant 201**. 5d.** Kraken 2 relative abundance in Participant 201.

To exclude the possibility that low-abundance or database-absent taxa were missed, we cross-validated MetaPhlAn4 profiles against Kraken2 for participant 201 (**Fig. 5c-d**). Consistent with prior benchmarking of these tools [26, 27, 33, 34, 36], the two pipelines showed strong concordance in diversity metrics and community composition (r = 0.76–0.92), with Kraken2 offering finer resolution within select genera (e.g., *Peptoniphilus*, *Streptococcus anginosus* group). The expanded taxonomic resolution revealed no additional candidate drivers of infection onset beyond those identified by MetaPhlAn4 (**Table S6–S7**).

### Antibiotic exposure, not catheter change, drives community instability and taxon abundance shifts

A formal longitudinal mixed-effects analysis was conducted across all nine participants, which identified antibiotic exposure — not catheter change — as the primary driver of community instability. Antibiotic exposure was consistently and significantly associated with overall community composition (PERMANOVA, Aitchison and Bray-Curtis, adjusted p = 0.006; **Fig. 6a**), and this signal remained robust when the most clinically extreme participant (201, two CAUTIs) and the most stable participant (104) were each excluded in leave-one-out tests (adjusted p ≤ 0.03 across all configurations; **Fig. S4**). In contrast, catheter change alone was not a consistent predictor of instability (adjusted p = 0.4 to >0.9), though an antibiotic-by-catheter interaction was detectable for Bray-Curtis (adjusted p = 0.016; **Fig. 6a**), suggesting catheter changes modulate, but do not independently drive, community disruption.

**Fig 6:**
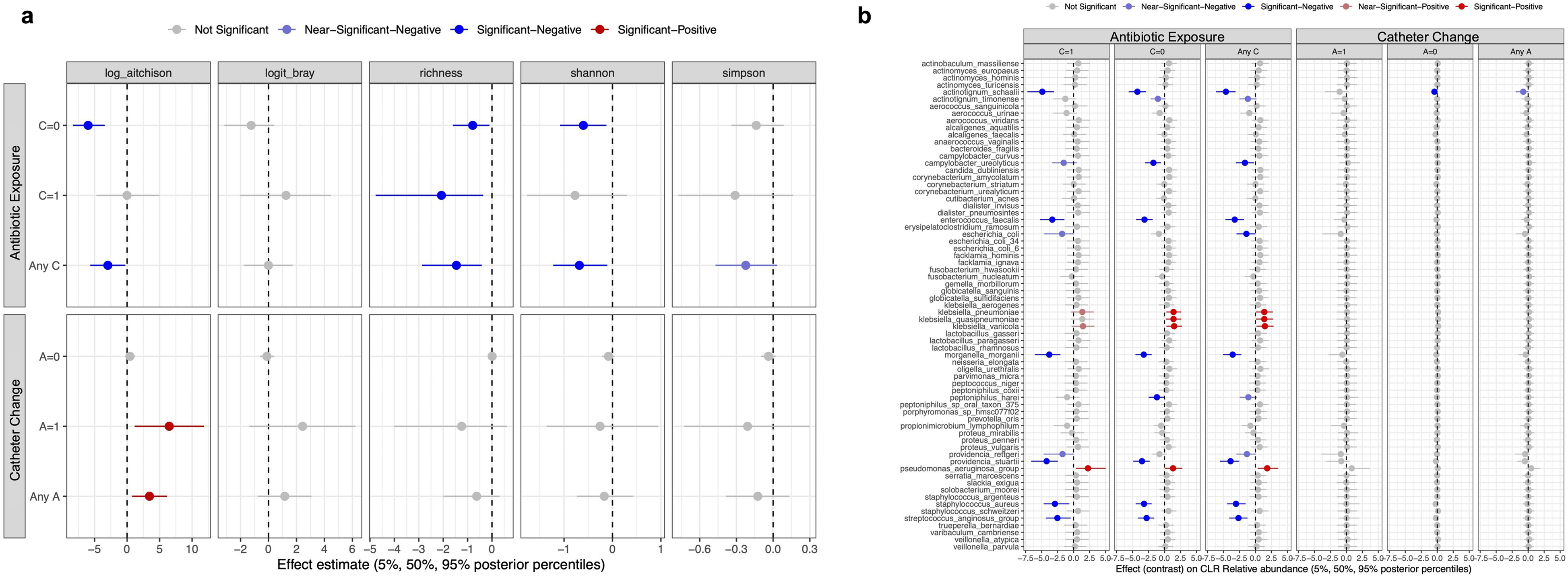
Antibiotic use and catheter change. 6a. Posterior estimates from five Bayesian linear mixed-effects models evaluating associations of antibiotic use (A), catheter change (C), and their interaction (A×C) with community-level outcomes across longitudinal urine metagenomes from 9 catheterized participants. Outcomes shown in columns are log-Aitchison distance between consecutive visits, logit-transformed Bray-Curtis dissimilarity, species richness, Shannon diversity, and Simpson index respectively between consecutive visits. Richness was defined as the number of taxa with relative abundance _≥_0.5%. **6b.** Species-level responses to antibiotic use and catheter change, and residual species-species association structure. Forest plot of posterior effect contrasts from a Bayesian Gaussian mixed-effects model of centered log-ratio (CLR) abundance for 69 taxa.

Species-level Bayesian modelling (**Fig. 6b**) across all 69 taxa revealed that this antibiotic effect was ecologically heterogenous rather than uniform community-wide loss. The model detected 15 significant or near-significant changes due to antibiotic exposure, regardless of catheter change status: four species tended to increase after antibiotic exposure (*Klebsiella pneumoniae, K. quasipneumoniae, K variicola,* and *Pseudomonas aeruginosa* group), while 11 decreased (*Actinotignum schaalii, A. timonense, C. ureolyticus, E. faecalis, E. coli, M. morganii, P. harei, P. rettgeri, P. stuartii, S. aureus,* and *S. anginosus* group). By contrast, catheter changes only produced a detectable effect on abundance of a single species (*Actinotignum schaalii*). These species-specific shifts remained robust when sensitivity analyses were repeated after excluding participants 201 and/or 104 (**Fig. S5**). Thus, antibiotic exposure perturbs the urinary community in a way that catheter changes do not. The species most consistently decreased by antibiotics were classical urinary opportunists (*E. faecalis, P. stuartii, M. morganii, S. aureus,* and *A. schaalii*), while species like *Klebsiella* and *Pseudomonas* increased after exposure either due to resistance to the antibiotic, lack of competition after loss of the opportunists, or some combination of both factors.

### Community-level co-occurrence analysis reveals linked species and mutually-exclusive clusters

The rich, longitudinal metagenomics datasets across multiple participants allowed us to examine microbial co-occurrences and determine whether there are associations among taxa that are reproducible across different individual participants. We first examined taxa co-occurrence, including whether co-occurring species experience similar abundance shifts after perturbations such as antibiotic exposure and catheter change. The co-occurrence matrices revealed 6 broad co-occurrence groups (mean correlation 0.46; 55 species pairs exceeded |r| > 0.30; labelled 1-6 in **Fig. 7a**, **Table 5**): 1) a <u>fastidious-anaerobe cluster</u> that represents the most internally-connected group in the matrix; 2) a <u>single pair</u> of *M. morganii* with *P. stuartii* (ρ = 0.43); 3) an <u>Enterobacterales cluster</u>; 4) a cluster of <u>urease-producers</u>; 5) <u>a Gram-</u> <u>negative/fastidious satellite cluster</u>; and 6) a <u>skin-associated cluster</u>. Beyond the six co-occurrence groups, two others stood out (labelled #, **Fig. 7a**): *S. anginosus* group and *Campylobacter ureolyticus* (ρ = 0.29), and a loose enteric/environmental Gram-negative group.

**Fig. 7.**
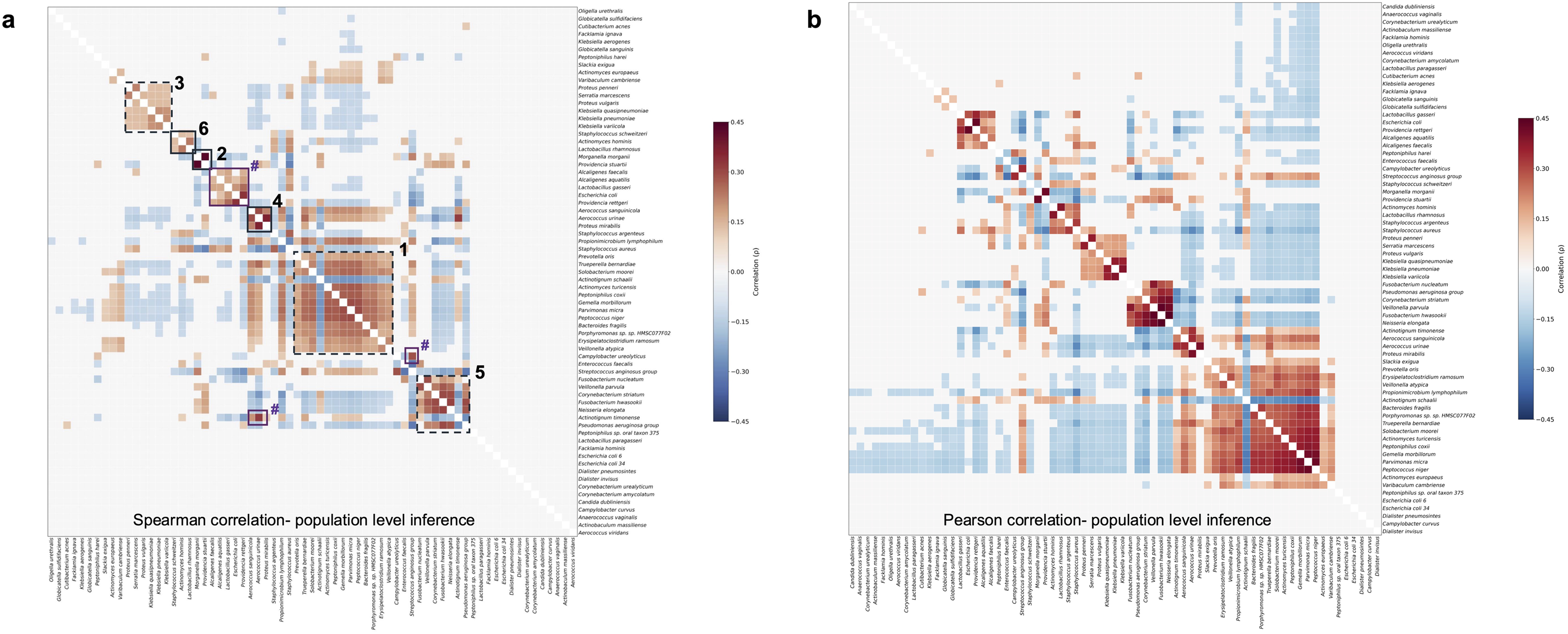
Correlation matrices from population-level inference. 7a. Spearman residual species-species association heatmaps derived from the Bayesian null mixed-effects model. **7b.** Pearson residual correlation matrices as a visualization cross-check.

**Table 5.**
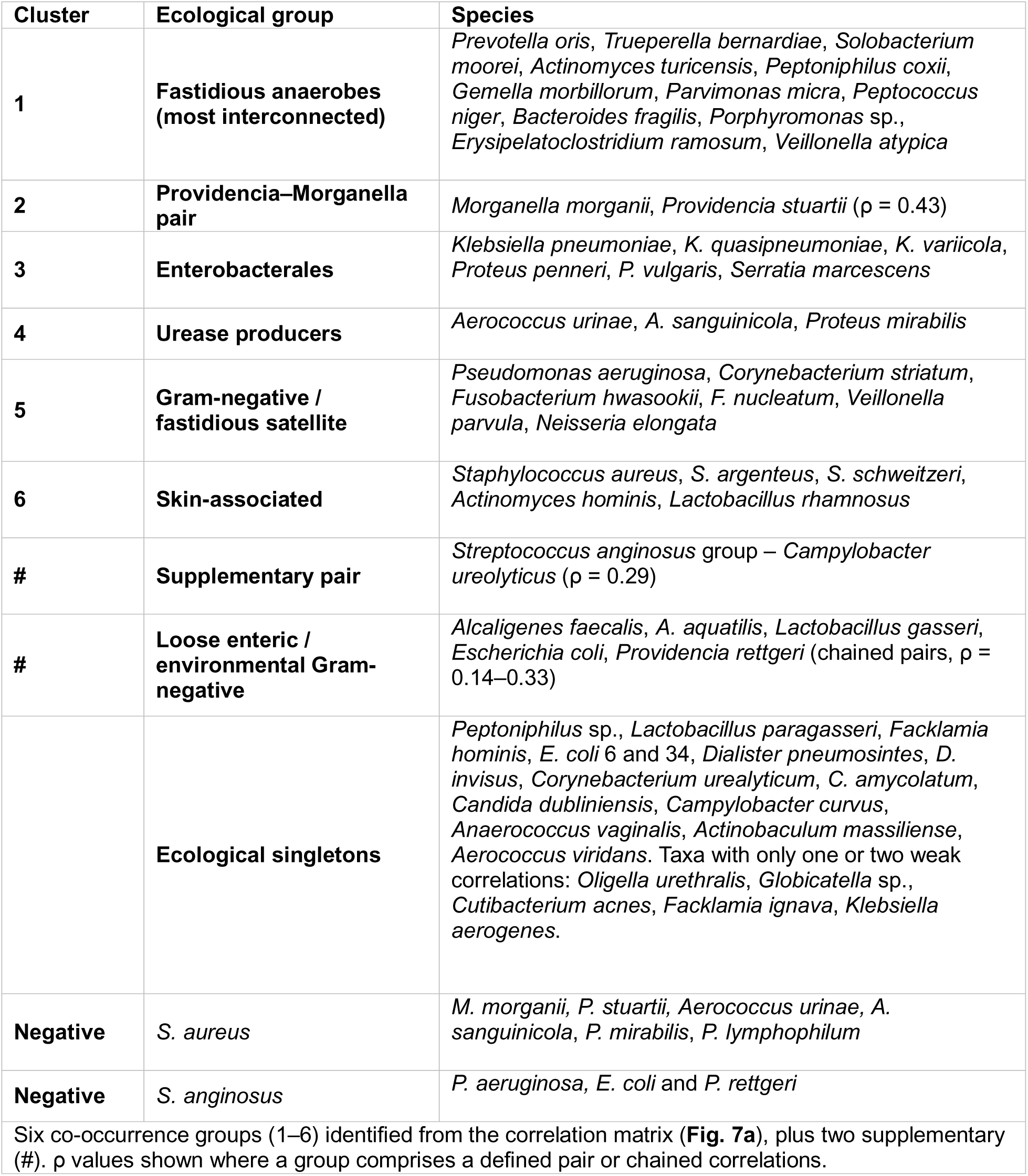
Co-occurrence of the catheter-associated urinary microbiome.

Negative correlations were almost as common as positive ones (47% positive among |ρ| > 0.10), with the strongest mutually-exclusive relationships involving *S. aureus* with cluster 2, urease producing cluster 4 (*Aerococcus urinae, A. sanguinicola*, and *P. mirabilis*), and *P. lymphophilum.* The next strongest negative correlations involved the *S. anginosus* group with cluster 5 (*P. aeruginosa)* and the Gram-negative group (*E. coli* and *P. rettgeri)*. Negative correlations also indicated mutual exclusivity where urine specimens enriched for one cluster, such as the Enterobacterales group (cluster 1), tended to be depleted of species in another fastidious-anaerobe group (cluster 1). *A. schaalii* also exhibited a strong negative correlation with all species in the fastidious-anaerobe group (cluster 1), while *A. timonense* opposed the Gram-negative/fastidious satellite group (cluster 5) but a positively correlated with *S. anginosus*. Additionally, approximately one-third of species behaved as ecological singletons with no co-variation with the rest of the community (**Table 5**). These species generally appeared and disappeared from individual urine samples without the co-occurrence or displacement of other species.

### Abundance shifts of key species may predict infection sign and symptom onset

Having characterized which species are antibiotic-responsive and which co-occur, we next asked whether abundance changes in any taxa or co-occurring groups correspond directly to infection sign and symptom onset by extending the co-occurrence and antibiotic-exposure observations to a possible clinical prediction framework. To do so, we fit logistic mixed-effects models linking species-level abundance to four CAUTI classification schemes: 1) potential CAUTI based on National Healthcare Safety Network (NHSN) criteria (the most stringent definition), 2) potential CAUTI based on NHSN criteria allowing for inclusion of highly polymicrobial urine cultures, 3) potential CAUTI based on Infectious Disease Society of America (IDSA) criteria considering only new-onset signs and symptoms, and 4) potential CAUTI based on IDSA criteria regardless of whether symptoms were new-onset (**Fig. 8, Fig. S6**). 24 species were identified with significant or near-significant associations with at least one definition, with 12 enriched in potentially symptomatic urines and 12 depleted (**Table 6**).

**Fig 8.**
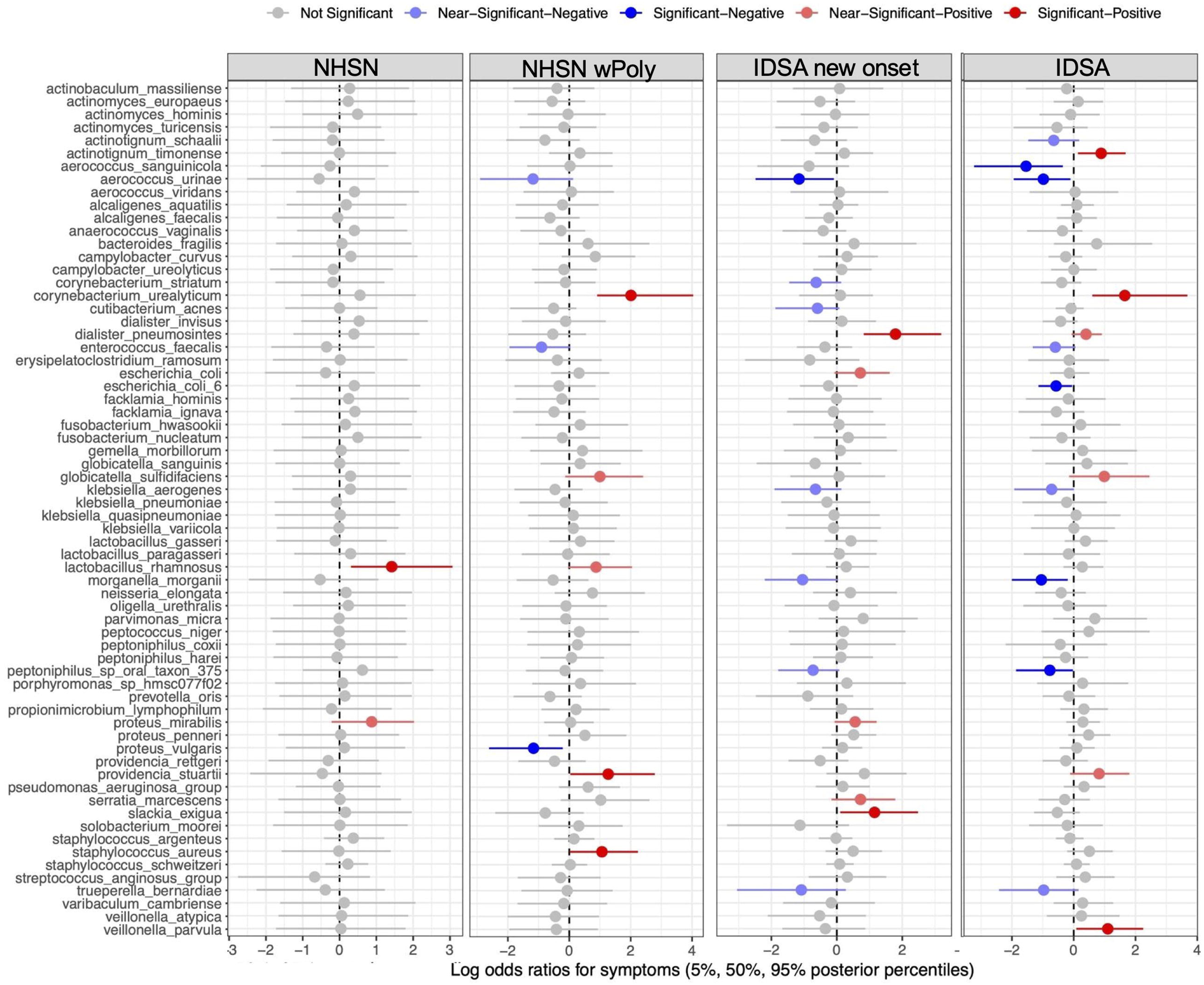
Standardized species CLR abundances associated with CAUTI/UTI outcome definitions with plot of posterior log-odds-ratio coefficients from Bayesian logistic mixed-effects models evaluating associations between standardized species CLR abundances and 4 binary clinical outcomes: potential CAUTI by IDSA criteria (IDSA), potential CAUTI by IDSA criteria restricted to new-onset symptoms (IDSA-new-onset), potential UTI by NHSN criteria (NHSN), and potential CAUTI by NHSN criteria allowing polymicrobial cultures (NHSN-wPoly). Each row is one species predictor.

**Table 6:**
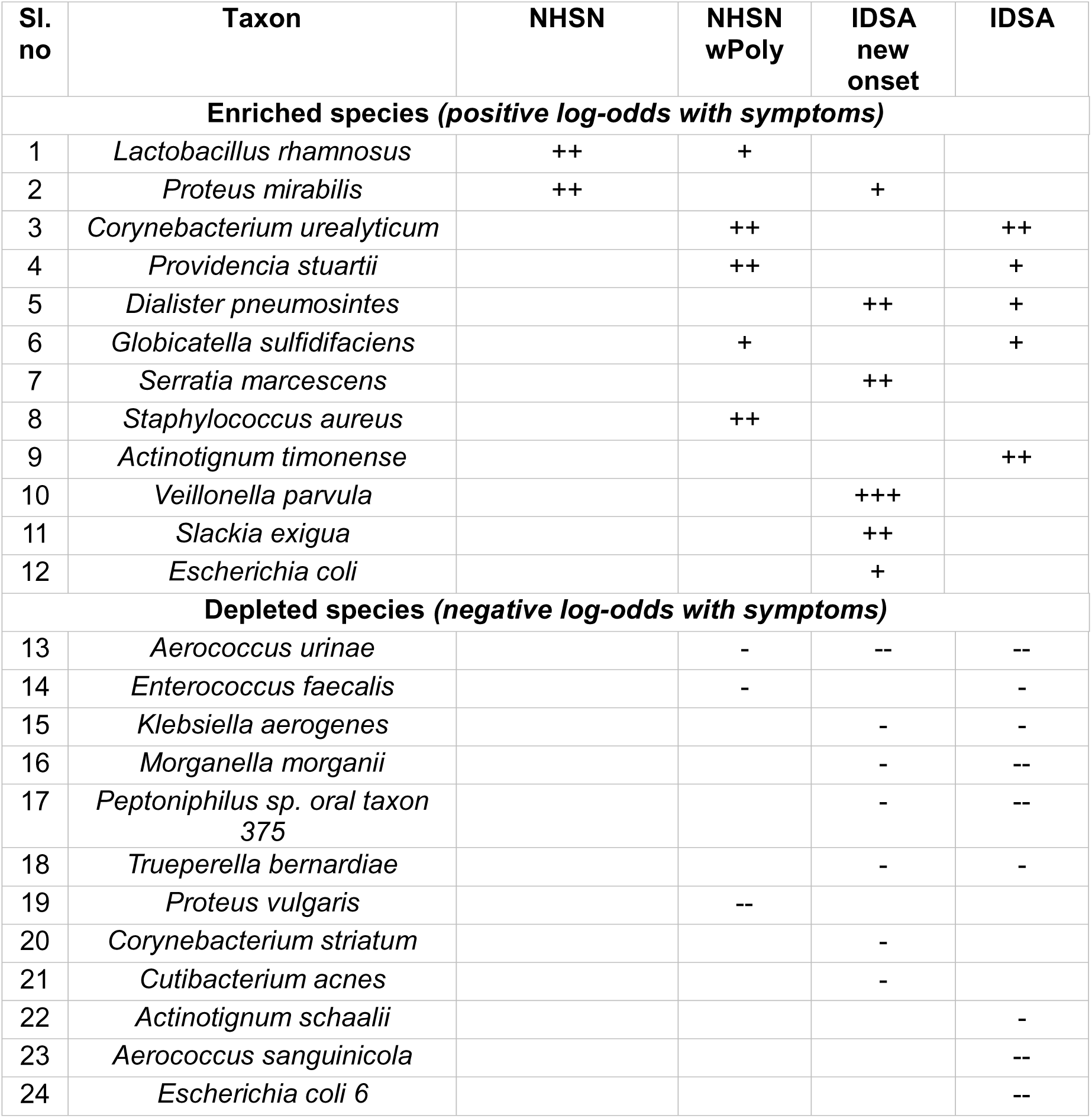
List of species for which abundance positively correlated (enriched) or negatively correlated (depleted) with CAUTI signs and symptoms.

Of the 12 species that were enriched during potentially symptomatic episodes, only 4 were depleted by antibiotics (*E. coli, P. stuartii, S. aureus,* and *A. timonense*), suggesting that treatment may fail to deplete clinically significant symptomatic species, especially those that would be missed by standard culture. It is also notable that only three of the enriched species met the threshold to be considered co-occurring: *S. aureus* has a slight positive association with *L. rhamnosus* (ρ = +0.20) and a weaker positive association with *S. marcescens* (ρ = +0.16). *S. aureus* abundance also has a negative correlation with two of the other enriched species: *P. stuartii* (ρ = -0.30) and *P. mirabilis* (ρ = -0.23). Hence, the species for which abundance was positively associated with infection signs and symptoms do not form a cohesive pro-disease community, suggesting that the abundances of these species may be independent predictors of symptomatic infection.

For the 12 species that were negatively correlated with infection signs and symptoms, their co-occurrence pattern was more complex. Three of the species have strong positive co-occurrences: *A. urinae* with *A. sanguinicola* (+0.29), *A. urinae* with *T. bernardiae* (+0.25), and *T. bernardiae* with *A. sanguinicola* (+0.22). Another species, *A. schaali,* also has weak positive associations with two other species from the list: *M. morganii* (+0.17) and *C. striatum* (+0.16). Thus, perturbations that decrease abundance of any one of these species may cause a corresponding decrease in the others. Furthermore, 8/12 species that were positively correlated with infection signs and symptoms exhibited negative cross-correlations with species that were negatively correlated with signs and symptoms. Thus, species that may have a protective effect for the host are interconnected and likely to decrease in abundance when symptom-associated species increase in abundance.

It is also notable that three of the species that were negatively correlated with symptoms were also depleted by antibiotics (*M. morganii, A. schaalii,* and *E. faecalis*), suggesting that antibiotic exposure can have the unintended consequence of depleting potential uropathogenic competitors. *M. morganii* abundance is negatively correlated with both *S. aureus* (ρ = −0.25) and *L. rhamnosus* (ρ = −0.16, modest), indicating that antibiotic suppression of *M. morganii* therefore removes a competitor of the *S. aureus*/*L. rhamnosus* axis that is positively correlated with symptoms. Similarly, *A. schaalii* abundance negatively correlates with *P. mirabilis* abundance (ρ = −0.21), suggesting that antibiotic suppression of *A. schaalii* can remove a competitor of the *P. mirabilis*/*A. timonense* axis that is positively correlated with symptoms. These networks are consistent with the idea that species such as *M. morganii* and *A. schaalii* may provide competitive resistance against symptom-associated organisms, with antibiotic exposure disrupting the ecological constraints that normally keep species such as *S. aureus* and *P. mirabilis* in check.

Based on the ecological framework of each participant, infection risk in long-term catheterized patients may also be more closely tied to the relative abundance of uropathogens versus commensals than to the mere culture-based presence of a pathogen. Participants 104 and 206 both carried a high uropathogen burden by standard culture (>10LJ CFU/ml at every visit, [5]), yet remained largely asymptomatic, with pathogens offset by a dominant, resilient backbone of oral commensals and anaerobes (**Table 7**). There observations point to a testable hypothesis that abundant commensals, coupled with species capable of modulating pathogenic potential (e.g., *M. morganii*) [37], may keep uropathogens in check and prevent the transition to symptomatic infection.

**Table 7.**
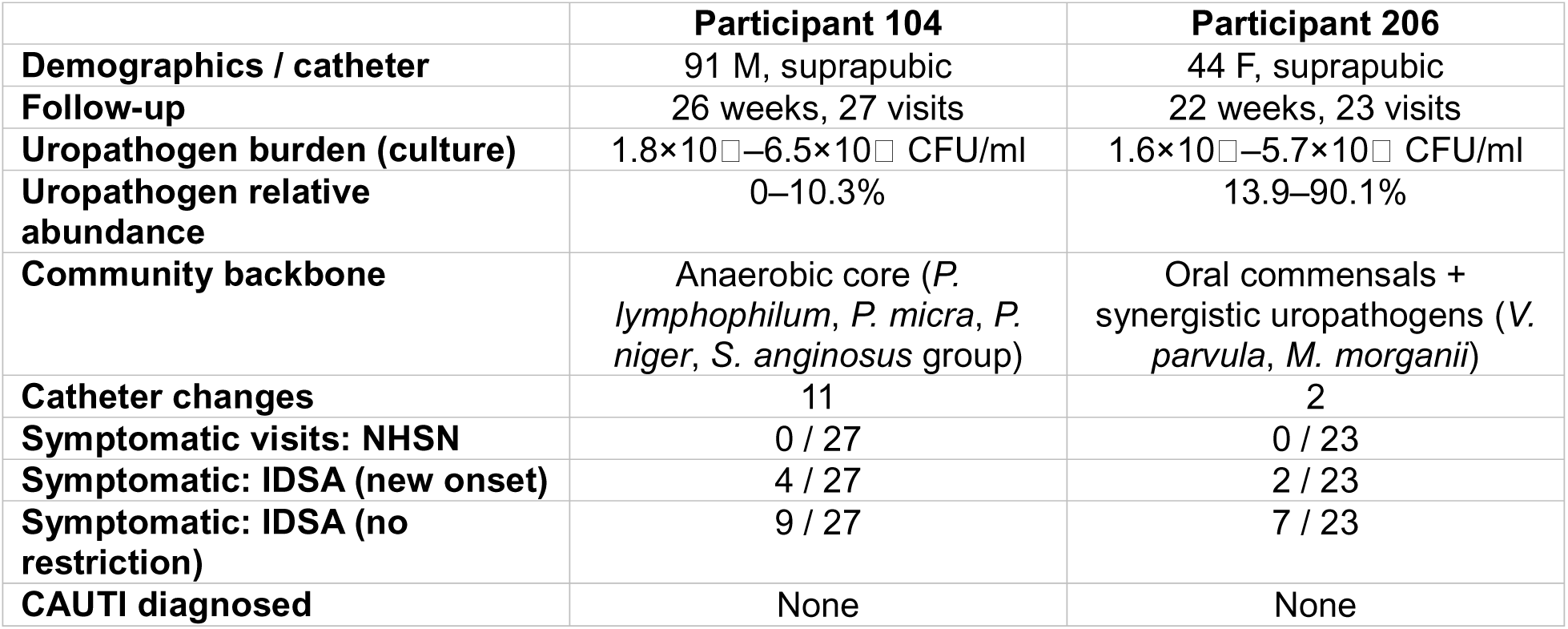
High culture-based uropathogen burden is decoupled from symptomatic infection in two stable participants.

## DISCUSSION

The long-term catheterized urinary tract has been characterized through the narrow lens of standard culture. Our longitudinal metagenomic data reveal a far more dynamic and compositionally complex ecosystem than previously described. Our findings demonstrate that the microbial composition of urine from individuals with long-term catheters is predominantly composed of established uropathogens (51.6% ± 33.5% of relative abundance across 198 samples in 9 participants). However, the abundance of additional species that would be missed by standard urine culture methods may critically shape transition risk from asymptomatic colonization to active infection. Pathogenic species versud commensals and oral microbes tend to co-occur and the abundance in these oral/commensal clusters are negatively associated with uropathogen cluster. Notably, of the 24 species that significantly correlated with the presence of CAUTI signs and symptoms, 11 cannot be detected by standard urine cultures. Thus, our data suggest that infection risk due to a high burden of uropathogens may be mitigated by the presence of specific commensal species and oral anaerobes, and this protective structure can be disturbed by antibiotic exposure.

In contrast to the healthy female urobiome, in which *Lactobacillus* species comprise 60-90% of the microbiota [8, 9, 38, 39], *Lactobacillus* species were rare in our catheterized population. Only 3/9 participants had *Lactobacillus* species (*L. gasseri, L. paragasseri, L. rhamnosus*). Of the 3 female participants in this cohort (102, 206, and 207), only 2 harbored *Lactobacillus* sp. and in very low abundance (0.09-6.3%), which may contribute to the high susceptibility to uropathogen overgrowth in female participants. However, a high abundance (0.05-47.38%) of *L. gasseri* was observed in male participant 106. This near absence of *Lactobacillus* in catheterized individuals and its negative association with the uropathogen cluster in our co-occurrence analysis is consistent with loss of pathogen colonization resistance in this population.

Our study also highlights *L. rhamnosus* as a potential opportunistic pathogen in this environment rather than a protective commensal. *L. rhamnosus* is widely known as a beneficial organism and a marker of healthy vaginal flora that exerts protective effects [40–43]. However, in our cohort of long-term catheterized individuals, high abundance of *L. rhamnosus* was significantly associated with potential symptomatic CAUTI under the NHSN definition (++), placing it among the symptom-enriched organisms in the dataset. Ecologically, *L. rhamnosus* was one of the few species in the matrix to show a stable, reproducible co-occurrence with *S. aureus* (ρ = +0.20). It also co-occurred with *A. hominis* (ρ = +0.23) and with the broader *S. aureus* complex (*S. argenteus,* ρ = +0.15), and was negatively correlated with the antibiotic-suppressed community moderator *M. morganii* (ρ = −0.16). Consequently, *L. rhamnosus* in catheterized urine may behave not as a vaginal-tract probiotic but rather as a skin-derived co-colonizer that travels alongside *S. aureus*. If correct, probiotic strategies built on the assumption that any *Lactobacillus* species is beneficial in the genitourinary tract may need to be reconsidered in this unique patient population.

The detection of oral species in 6/9 participants suggests potential oral-urinary bacterial translocation, including the presence of periodontal pathogens such as *Fusobacterium nucleatum* (4/9), *Streptococcus anginosus* group (4/9), and *Parvimonas micra* (2/9). These organisms are co-associated with polymicrobial abscesses and periodontal disease, raising questions about route of colonization in the urinary tract [28, 44–47]. In our co-occurrence analysis, these oral-associated taxa formed a distinct co-occurrence cluster that was mutually exclusive with the uropathogen-associated cluster, suggesting that oral species may occupy an ecological niche antagonistic to uropathogen colonization. However, whether this reflects active competitive exclusion in common environmental conditions (e.g., pH, oxygen need) cannot be determined from the current data.

The presence of commensal species (30.7%), particularly urogenital commensals such as *Actinotignum schaalii* (7/9 participants) and *Aerococcus urinae* (4/9), supports the emerging concept of a core urinary microbiome as residents of the urinary tract, particularly in elderly and catheterized individuals [32, 38, 39]. *A. schaalii* is recognized as a common colonizer of the elderly urinary tract and has been associated with urinary tract infections in immunocompromised patients [48], and it was highly prevalent in our cohort (89% of participants). As *A. schaalii* was negatively-correlated with CAUTI signs and symptoms as well as with uropathogens such as *P. mirabilis,* our data suggest that this species may act as a beneficial commensal in the catheterized urinary tract.

The negative correlation of *M. morganii* with CAUTI signs and symptoms is noteworthy since this species is considered an opportunistic pathogen of the urinary tract but can also play a modulatory role [37, 49]. In our cohort, *M. morganii* was suppressed by antibiotic exposure (β = −3.6, robust across all four sensitivity analyses). It co-occurred most closely with the symptom-associated species *P. stuartii* (ρ = +0.43) but was negatively correlated with both anchors of the *S. aureus*-associated symptomatic axis: *S. aureus* itself (ρ = −0.25) and *L. rhamnosus* (ρ = −0.16). Taken together, these patterns suggest that *M. morganii*, despite its conventional classification as an opportunistic uropathogen, may occupy a community-stabilizing role in the catheterized bladder of asymptomatic carriers. Its presence may constrain the symptom-associated *S. aureus* and *L. rhamnosus*, and its depletion following antibiotic exposure may coincide with the release of that constraint. This is consistent with our prior observations in which *M. morganii* played a potential role in decreased infection severity in part by dampening urease activity and cytotoxicity driven by other species [37, 50]. *Klebsiella aerogenes* also secretes urease-dampening compounds and has been hypothesized to similarly play a protective, disease-dampening role as *M. morganii* [50], which is supported by the negative-correlation of *K. aerogenes* abundance with infection signs and symptoms in our study. The broader implication of these observations is that organisms catalogued as opportunistic uropathogens based on isolated culture-positive episodes may behave very differently in the context of a stable polymicrobial community, and that antibiotic stewardship in catheterized patients should consider not only direct pathogen suppression but the loss of community moderators like *M. morganii* and *K. aerogenes* that may actually help maintain asymptomatic bacteriuria.

Antibiotic exposures were heterogeneous in indication: only 3 (Participants 101, 201) targeted the urinary tract, while the rest treated non-urinary infections. This distinction is important to interpret antibiotic-driven changes reflect broad spectrum antibiotic exposure rather than only UTI-specific effects. Participant 203 exemplifies this where 5 antibiotic courses for a persistent skin infection repeatedly restructured the urinary microbiota, driving dominance of resistant *P. aeruginosa*. Each cessation opened an ecological vacuum filled by *S. aureus* and *E. faecalis*, until the next course cleared them and *P. aeruginosa* reclaimed dominance. Though never diagnosed with CAUTI, this participant had 7/19 visits meeting some symptomatic threshold (1 by NHSN, 4–7 by IDSA depending on new-onset criteria), 4 of which followed antibiotic exposure (weeks 4, 5, 6, 20). Whether symptom onset reflects microbial dysbiosis, antibiotic side effects, or the underlying skin infection cannot be determined, but this case suggests that antibiotic exposure even when not urinary-directed can act as a tipping point toward infection in participants with low richness/diversity or high uropathogen abundance.

Overall, our findings suggest a possible ecological framework for CAUTI risk management. A high-diversity, stable profile in an asymptomatic patient may argue against antibiotic intervention, whereas a low-diversity, pathogen-dominated profile lacking protective species could justify targeted treatment. Participants with high commensal burden typically remained asymptomatic despite harboring recognized uropathogens, while low-diversity communities were more ecologically vulnerable. This is supported by studies where higher microbial diversity is associated with lower UTI incidence [51–53]. Similarly, recent studies on spinal cord injured catheterized individuals, and non-catheterized recurrent UTI studies indicated decreased diversity to be linked with high uropathogens [31, 54]. Future studies with larger cohorts, standardized clinical endpoints, and longitudinal sampling designs can be tested to improve the prediction of CAUTI onset beyond conventional microbiological criteria.

Formal cluster analyses were not performed due to the small sample size; the ecological phenotypes described here represent descriptive patterns requiring confirmation in larger cohorts. The inter-individual variability observed is consistent with personalized within-host signatures [14, 17, 55–57], likely shaped by host immune status, duration and type of catheterization, and antibiotic exposure history. With only 9 participants, larger interventional studies are needed to confirm causal links between ecological dynamics and clinical outcomes. All longitudinal Bayesian analyses should be interpreted as exploratory, with uncertainty summarized by posterior intervals and lfsr rather than proof of causation. Age was described clinically but not modeled, so age-adjusted inference is not possible. Finally, the oral/commensal correlation while statistically significant, reflects published primary habitat and not behavior in urobiome ecology [28]. Future work incorporating strain level analysis, host immune profiling, metabolomics, and detailed clinical annotations will provide a more integrated understanding of the host-microbe interface driving symptom onset.

## CONCLUSIONS

The catheterized urinary tract is a dynamic ecosystem governed by stability, resilience, and collapse, with the catheterized urinary tract ranging from robust, colonization-resistant communities to fragile, pathogen-dominated states. Unlike the *Lactobacillus*-dominated urobiome, catheterized individuals harbor *Lactobacillus*-poor communities enriched in *Providencia*, *M. morganii*, and oral pathogens. Recognizing these phenotypes lays the foundation for precision CAUTI management guided by infection ecology to improve diagnosis, refine antibiotic use, and better prevent one of healthcare’s most recalcitrant infections.

## Availability of data and materials

All data generated or analysed during this study are included in this published article and its supplementary information files. The sequencing data generated in this study is publicly available under BioProject: PRJNA1466380. All scripts used in this study can be found in Additional File 1.

## Authors’ Contributions

CEA and VSC conceptualized the study, acquired funding and provided resources. ALB tested all the DNA extraction kits for optimization and isolated DNA from all 198 samples. EMN performed DNA sequencing. ND and EMN carried out the metagenomic analysis. ND wrote the manuscript and prepared the figures. SC performed the longitudinal statistical analyses and prepared the corresponding figures. All authors critically reviewed and edited the manuscript and approved the final version for submission.

## Supporting information

Supplemental Scripts

## Acknowledgements

Not Applicable

## References

1. Armbruster, C.E., H.L.T. Mobley, and M.M. Pearson, Pathogenesis of Proteus mirabilis Infection. EcoSal Plus, 2018. 8(1).

2. Amer, M.A., et al., Silicone Foley catheters impregnated with microbial indole derivatives inhibit crystalline biofilm formation by Proteus mirabilis. Front Cell Infect Microbiol, 2022. 12: p. 1010625.

3. Armbruster, C.E., et al., How Often Do Clinically Diagnosed Catheter-Associated Urinary Tract Infections in Nursing Homes Meet Standardized Criteria? J Am Geriatr Soc, 2017. 65(2): p. 395–401.

4. Gaston, J.R., et al., Polymicrobial interactions in the urinary tract: is the enemy of my enemy my friend? Infect Immun, 2021.

5. Armbruster, C.E., et al., Prospective assessment of catheter-associated bacteriuria clinical presentation, epidemiology, and colonization dynamics in nursing home residents. JCI Insight, 2021. 6(19).

6. Magill, S.S., et al., Changes in Prevalence of Health Care-Associated Infections in U.S. Hospitals. N Engl J Med, 2018. 379(18): p. 1732–1744.

7. Scott, R.I., The Direct Medical Costs of Healthcare-Associated Infections in U.S. Hospitals and the Benefits of Prevention. Centers for Disease Control and Prevention. . 2009.

8. Hilt, E.E., et al., Urine is not sterile: use of enhanced urine culture techniques to detect resident bacterial flora in the adult female bladder. J Clin Microbiol, 2014. 52(3): p. 871–6.

9. Thomas-White, K., et al., Culturing of female bladder bacteria reveals an interconnected urogenital microbiota. Nat Commun, 2018. 9(1): p. 1557.

10. Nye, T.M., et al., Microbial co-occurrences on catheters from long-term catheterized patients. Nat Commun, 2024. 15(1): p. 61.

11. Jacobsen, S.M., et al., Complicated catheter-associated urinary tract infections due to Escherichia coli and Proteus mirabilis. Clin Microbiol Rev, 2008. 21(1): p. 26–59.

12. Duran Ramirez, J.M., et al., Semi-Quantitative Assay to Measure Urease Activity by Urinary Catheter-Associated Uropathogens. Front Cell Infect Microbiol, 2022. 12: p. 859093.

13. Didelot, X., et al., Within-host evolution of bacterial pathogens. Nat Rev Microbiol, 2016. 14(3): p. 150–62.

14. Thanert, R., et al., Persisting uropathogenic Escherichia coli lineages show signatures of niche-specific within-host adaptation mediated by mobile genetic elements. Cell Host Microbe, 2022. 30(7): p. 1034–1047 e6.

15. Young, B.C., et al., Evolutionary dynamics of Staphylococcus aureus during progression from carriage to disease. Proc Natl Acad Sci U S A, 2012. 109(12): p. 4550–5.

16. Bentley, S.D. and J. Parkhill, Genomic perspectives on the evolution and spread of bacterial pathogens. Proc Biol Sci, 2015. 282(1821): p. 20150488.

17. Sibale, L.L., et al., Within-host genetic diversity of pneumococcal serotype 3 during one-year prolonged carriage in a healthy adult. Nat Commun, 2025. 16(1): p. 8920.

18. Heravi, F.S., et al., Host DNA depletion efficiency of microbiome DNA enrichment methods in infected tissue samples. J Microbiol Methods, 2020. 170: p. 105856.

19. Davis, A., et al., Improved yield and accuracy for DNA extraction in microbiome studies with variation in microbial biomass. Biotechniques, 2019. 66(6): p. 285–289.

20. Shaffer, J.P., et al., A comparison of six DNA extraction protocols for 16S, ITS and shotgun metagenomic sequencing of microbial communities. Biotechniques, 2022. 73(1): p. 34–46.

21. Nelson, M.T., et al., Human and Extracellular DNA Depletion for Metagenomic Analysis of Complex Clinical Infection Samples Yields Optimized Viable Microbiome Profiles. Cell Rep, 2019. 26(8): p. 2227–2240 e5.

22. Harris, P.A., et al., The REDCap Mobile Application: a data collection platform for research in regions or situations with internet scarcity. JAMIA Open, 2021. 4(3): p. ooab078.

23. Harris, P.A., et al., The REDCap consortium: Building an international community of software platform partners. J Biomed Inform, 2019. 95: p. 103208.

24. Harris, P.A., et al., Research electronic data capture (REDCap)--a metadata-driven methodology and workflow process for providing translational research informatics support. J Biomed Inform, 2009. 42(2): p. 377–81.

25. Franzosa, E.A., et al., Species-level functional profiling of metagenomes and metatranscriptomes. Nat Methods, 2018. 15(11): p. 962–968.

26. Beghini, F., et al., Integrating taxonomic, functional, and strain-level profiling of diverse microbial communities with bioBakery 3. Elife, 2021. 10.

27. Segata, N., et al., Metagenomic microbial community profiling using unique clade-specific marker genes. Nat Methods, 2012. 9(8): p. 811–4.

28. Chen, T., et al., The Human Oral Microbiome Database: a web accessible resource for investigating oral microbe taxonomic and genomic information. Database (Oxford), 2010. 2010: p. baq013.

29. Sohngen, C., et al., BacDive--the Bacterial Diversity Metadatabase. Nucleic Acids Res, 2014. 42(Database issue): p. D592–9.

30. Hooton, T.M., et al., Diagnosis, Prevention, and Treatment of Catheter-Associated Urinary Tract Infection in Adults: 2009 International Clinical Practice Guidelines from the Infectious Diseases Society of America. Clinical Infectious Diseases, 2010. 50(5): p. 625–663.

31. Worby, C.J., et al., Longitudinal multi-omics analyses link gut microbiome dysbiosis with recurrent urinary tract infections in women. Nat Microbiol, 2022. 7(5): p. 630–639.

32. Wolfe, A.J., et al., Evidence of uncultivated bacteria in the adult female bladder. J Clin Microbiol, 2012. 50(4): p. 1376–83.

33. Lu, J., et al., Metagenome analysis using the Kraken software suite. Nat Protoc, 2022. 17(12): p. 2815–2839.

34. Wood, D.E., J. Lu, and B. Langmead, Improved metagenomic analysis with Kraken 2. Genome Biol, 2019. 20(1): p. 257.

35. Dudeck, M.A., et al., National Healthcare Safety Network report, data summary for 2013, Device-associated Module. Am J Infect Control, 2015. 43(3): p. 206–21.

36. Davis, M.P., et al., Kraken: a set of tools for quality control and analysis of high-throughput sequence data. Methods, 2013. 63(1): p. 41–9.

37. Learman, B.S., et al., A Rare Opportunist, Morganella morganii, Decreases Severity of Polymicrobial Catheter-Associated Urinary Tract Infection. Infect Immun, 2019. 88(1).

38. Pearce, M.M., et al., The female urinary microbiome: a comparison of women with and without urgency urinary incontinence. mBio, 2014. 5(4): p. e01283–14.

39. Pearce, M.M., et al., The female urinary microbiome in urgency urinary incontinence. Am J Obstet Gynecol, 2015. 213(3): p. 347 e1–11.

40. Tas, G.G. and L. Sati, Probiotic Lactobacillus rhamnosus species: considerations for female reproduction and offspring health. J Assist Reprod Genet, 2024. 41(10): p. 2585–2605.

41. Gardiner, G.E., et al., Persistence of Lactobacillus fermentum RC-14 and Lactobacillus rhamnosus GR-1 but not L. rhamnosus GG in the human vagina as demonstrated by randomly amplified polymorphic DNA. Clin Diagn Lab Immunol, 2002. 9(1): p. 92–6.

42. Pascual, L.M., et al., Lactobacillus rhamnosus L60, a potential probiotic isolated from the human vagina. J Gen Appl Microbiol, 2008. 54(3): p. 141–8.

43. Almurad, B., et al., Protective Effects of Lactobacillus rhamnosus GG Against Methotrexate-Induced Oxidative Renal Toxicity. Probiotics Antimicrob Proteins, 2026.

44. Chen, Y., et al., More Than Just a Periodontal Pathogen -the Research Progress on Fusobacterium nucleatum. Front Cell Infect Microbiol, 2022. 12: p. 815318.

45. Krishnan, K., T. Chen, and B.J. Paster, A practical guide to the oral microbiome and its relation to health and disease. Oral Dis, 2017. 23(3): p. 276–286.

46. Schmalz, G., et al., Dental and periodontal health, and microbiological and salivary conditions in patients with or without diabetes undergoing haemodialysis. Int Dent J, 2017. 67(3): p. 186–193.

47. Dugdale, C.M., et al., Cavernosal Abscess due to Streptococcus Anginosus: A Case Report and Comprehensive Review of the Literature. Curr Urol, 2013. 7(1): p. 51–6.

48. Lotte, R., L. Lotte, and R. Ruimy, Actinotignum schaalii (formerly Actinobaculum schaalii): a newly recognized pathogen-review of the literature. Clin Microbiol Infect, 2016. 22(1): p. 28–36.

49. Liu, H., et al., Morganella morganii, a non-negligent opportunistic pathogen. Int J Infect Dis, 2016. 50: p. 10–7.

50. Guterman, L.B., et al., Harnessing microbial-derived metabolites in the urinary tract to prevent infection induced catheter encrustation. Nature Communications, 2025. 16(1): p. 9678.

51. Moustafa, A., et al., Microbial metagenome of urinary tract infection. Sci Rep, 2018. 8(1): p. 4333.

52. Horwitz, D., et al., Decreased microbiota diversity associated with urinary tract infection in a trial of bacterial interference. J Infect, 2015. 71(3): p. 358–367.

53. Fouts, D.E., et al., Integrated next-generation sequencing of 16S rDNA and metaproteomics differentiate the healthy urine microbiome from asymptomatic bacteriuria in neuropathic bladder associated with spinal cord injury. . Journal of translational medicine, 1012. 10(174).

54. Hoque, M.M., et al., Prediction of symptomatic and asymptomatic bacteriuria in spinal cord injury patients using machine learning. Microbiome, 2025. 13(1): p. 246.

55. Amunugama, K., D.P. Pike, and D.A. Ford, E. coli strain-dependent lipid alterations in cocultures with endothelial cells and neutrophils modeling sepsis. Front Physiol, 2022. 13: p. 980460.

56. Guerillot, R., et al., Unstable chromosome rearrangements in Staphylococcus aureus cause phenotype switching associated with persistent infections. Proc Natl Acad Sci U S A, 2019. 116(40): p. 20135–20140.

57. Neugent, M.L., et al., Urinary biochemical ecology reveals microbiome-metabolite interactions and metabolic markers of recurrent urinary tract infection. NPJ Biofilms Microbiomes, 2025. 11(1): p. 216.

