## Supplemental Figures for "Optimized Urine Metagenomic Methods Reveal Longitudinal Microbial Community Dynamics and Predictors of Transition from Asymptomatic Colonization to CAUTI"

### Supplementary Figures

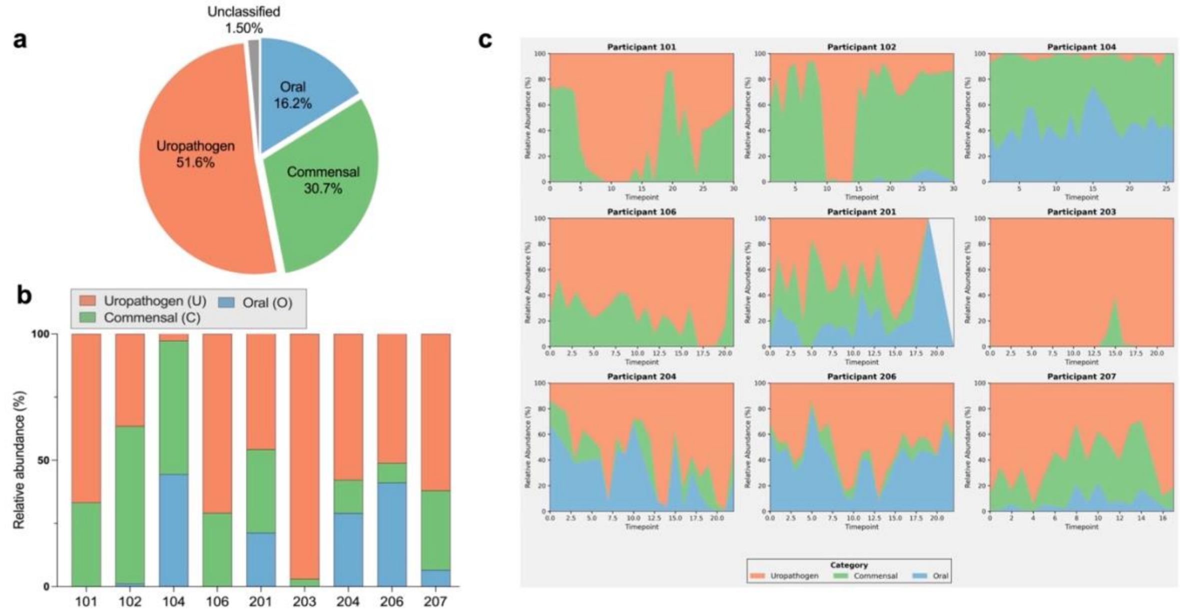


| N | 27 | 26 | 26 | 22 | 19 | 19 | 21 | 23 | 17 |
|---|---|---|---|---|---|---|---|---|---|
| Time range | 0-30 | 1-30 | 1-26 | 0-21 | 0-22 | 0-22 | 0-22 | 0-22 | 0-17 |
| Dominant | U | C | C | U | U | U | U | U | U |

**Fig. S1**. **Classification of microbial species into uropathogens, commensal microbes, and oral microbes. S1a**. Percentage of classified species. **S1b.** Relative abundance percentage of uropathogens, oral, and commensal microbes across 9 participants across N samples. 7/9 participants were dominated by uropathogens while 2/9 participants has commensal microbes. **S1c.** Relative abundance (%) of uropathogens, commensal, and oral microbes across the 9 study participants at all timepoints.

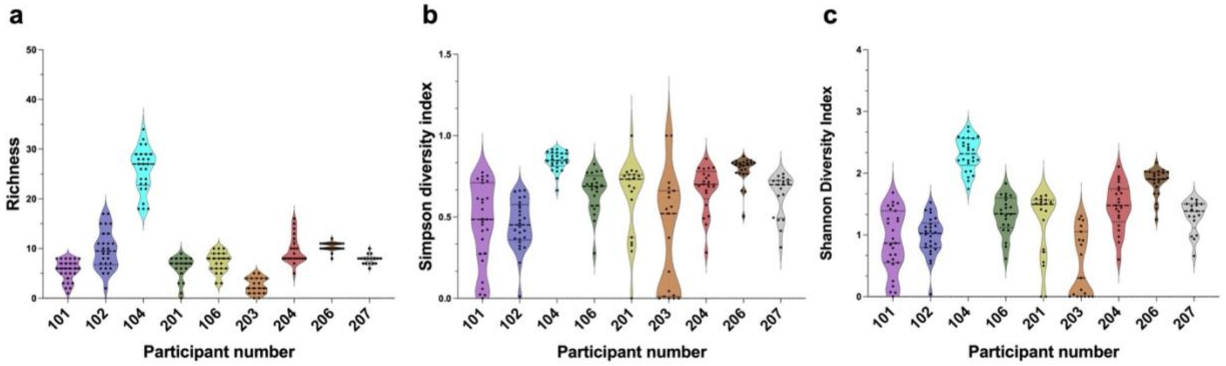


**Fig. S2. Inter-individual differences in microbial diversity across participants.** Distribution of **S2a.** species richness, **S2b.** Shannon diversity, and **S2c.** Simpson diversity across all participants. Statistical testing confirmed strong inter-individual differences for all three metrics (Kruskal–Wallis: richness = 141.77, $p < 0.001$; Shannon = 129.50, $p < 0.001$; Simpson = 102.71, $p < 0.001$). Participant 104 exhibited significantly higher diversity across all metrics. Participant 203 showed the lowest richness and diversity, reflecting a community dominated by few taxa.

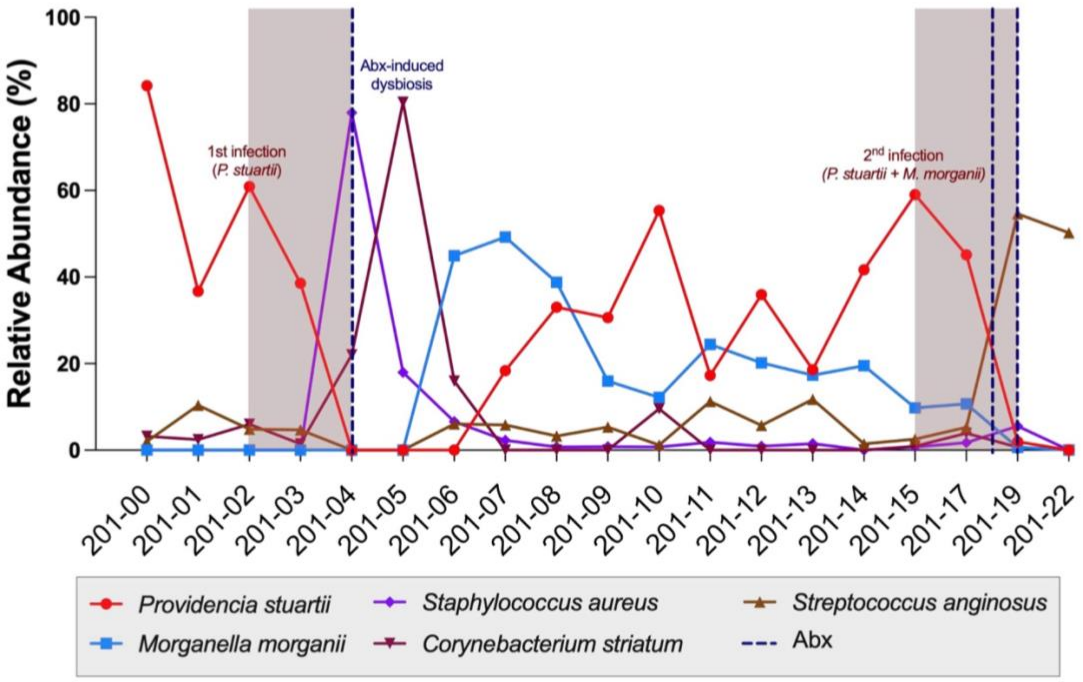


**Fig. S3:** Relative abundance (%) of uropathogens over time and infection cycles Participant 201.

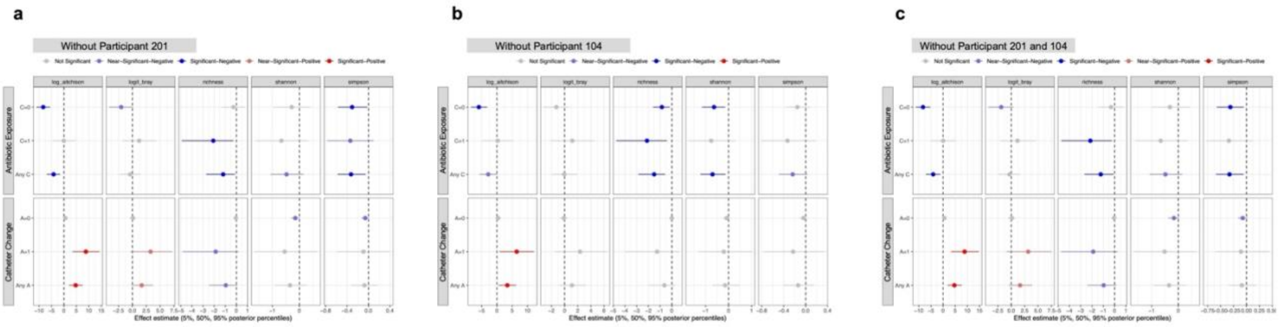


**Fig. S4: Leave-one-out sensitivity analysis for community-level exposure effects.** Sensitivity analyses after excluding participant **S4a.** 201, **S4b.** participant 104, or **S4c.** both participants. Participant 201 was the most clinically complex participant, whereas participant 104 was the highest-diversity ecological outlier.

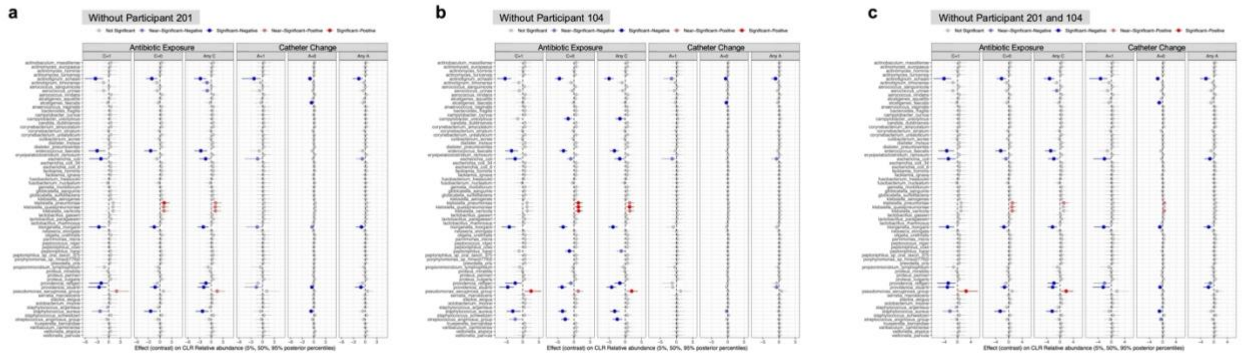


**Fig.S5: Full species-level contrast set and leave-one-out sensitivity analysis.** Species-level CLR effect contrasts from the Bayesian mixed-effects model, showing all six conditional and marginal exposure contrasts: antibiotic effects at C=0 and C=1 and averaged over catheter status (Any C), and catheter-change effects at A=0 and A=1 and averaged over antibiotic status (Any A).

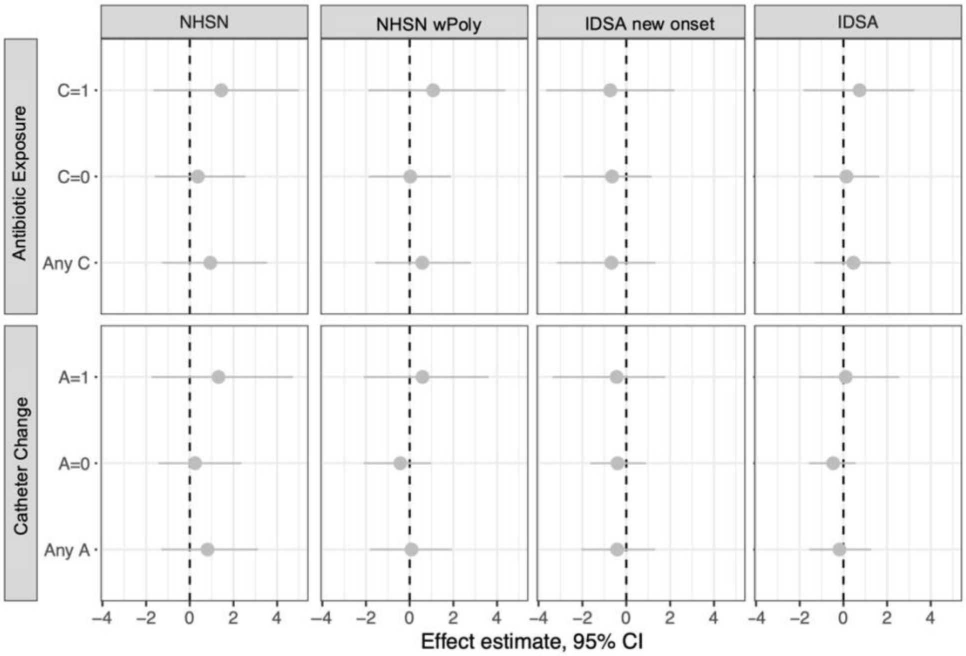


**Fig.S6: PERMANOVA sensitivity analysis for community-level exposure effects.** PERMANOVA results testing associations of antibiotic use and catheter change with between-sample community dissimilarity, reported as a non-Bayesian cross-check.

### Supplementary Note A

<u>*Participant 102.*</u> This 61-year-old female with a Foley catheter exhibited a moderate to highly diverse community structure in the first few weeks of the study (S=8 at baseline, S=15 at week 1, and H′ ~1.36). Microbial composition was largely stable over 30 weeks of follow-up and unaffected by catheter changes. Specifically, catheter changes occurred within the week before study visits 3, 7, 9, 11, 14, 20, 25, and 30, but did not alter $\alpha$-diversity (Kruskal-Wallis test, $p = 0.2660$) or change microbial composition (**Fig. 2a**). The participant did not have any known exposure to antibiotics during follow-up and was not diagnosed with CAUTI at any point.

From the baseline visit to week 9, the microbial community was consistently dominated by *Aerococcus urinae*. However, between weeks 10-14, flanked by 2 catheter changes at weeks 11 and 14, there was a transient bloom of *Enterococcus faecalis* followed by *Proteus mirabilis*. Both taxa were present in the weeks preceding the bloom, indicating an expansion from the resident microflora rather than a new introduction. The bloom coincided with temporary displacement of the dominant *A. urinae,* but the community quickly reset to the original structure by week 15 (**Fig. 2b**).

<u>*Participant 106.*</u> The microbial diversity in this 61-year-old male with a suprapubic catheter exhibited dynamic instability, with periodic changes in relative richness and dramatic $\alpha$-diversity loss. H′ ranged from 0.61-1.83, D from 0.28 to 0.83, and S varied from 3 to 10 observed species. The community maintained a moderate diversity in the first 9 weeks (H′>1.34) and was resilient to the first several catheter changes (weeks 1, 4, 9) with $\alpha$-diversity recovering and increasing within one week after the catheter change.

However, the catheter change at week 17 coincided with collapsed community diversity, with H′ dropping 45% to 0.81 by week 19. At the final timepoint (week 21) there was another collapse of the community structure with H′=0.61 and D=0.21 despite maintained richness (7 species) (**Fig. 2c**).

Two gram-positive species (*Staphylococcus aureus and E. faecalis)*, along with two Enterobacterales (*Escherichia coli* and *Klebsiella pneumoniae*) consistently dominated from baseline to week 16. *Lactobacillus gasseri* was also detected as a major recurring member, which is notable as 106 was the only participant for which this common gut and genitourinary commensal was detected in catheterized urine samples. *P. mirabilis* first appeared in low abundance at week 15 and then bloomed following the week 17 catheter change, corresponding to the collapse in α-diversity at week 18. When the *P. mirabilis* bloom receded after week 19, it was not replaced by the original composition but was succeeded by a new dominant species (*A. urinae)* at the final timepoint (week 21), with a shift to a low-diversity community (**Fig. 2d**).

Participant 204. The microbial diversity in this 51-year-old male with a suprapubic catheter demonstrated two periods of distinct community profiles. Prior to a catheter change at week 13, the microbial community exhibited relatively stable richness and α-diversity despite a transient decrease in the Shannon index from 1.58 at week 5 to 0.60 at week 6. Following a catheter change at week 13, a clear shift in community structure occurred. Richness increased from 10 at week 13 to 16 at week 14, and both Shannon and Simpson indices showed a sustained rise beginning at week 14, indicating a transition toward a more diverse and even microbial community. By week 22, diversity reached its peak, with a H' of 2.11 and a D of 0.86, reflecting the highest S of 14 taxa and evenness during the monitoring period (**Fig. 2e**).

During the initial period between baseline and week 12, *Streptococcus anginosus* group dominated the community. This dominant population co-occurred with a dynamic, polymicrobial background of recognized uropathogens, including *Morganella morganii*, *Alcaligenes faecalis*, *Serratia marcescens*, and *S. aureus*. A transient bloom of *S. marcescens* occurred at week 7 and resolved by week 8. The catheter change at week 13 corresponded to a major ecological shift: the previously dominant *S. anginosus* group was displaced by a newly acquired population of *K. pneumoniae*, a classic nosocomial uropathogen. From week 15 onward, *K. pneumoniae* remained the dominant taxon, though its abundance fluctuated with other organisms, including the persistent *M. morganii*. This second stable community also included several new species acquired at week 13, namely *Klebsiella variicola*, *Klebsiella quasipneumoniae*, and *Campylobacter ureolyticus*. (**Fig. 2f**).

<u>Participant 104</u>: This 91-year-old male with a suprapubic catheter had a very high diversity at baseline (S=27 and H′ ~2.21) and it remained consistently high throughout 26 weeks of follow-up, with S ranging between 18-38 and H′ ~1.7-2.7. Despite 11 catheter changes during the study, no significant impact on $\alpha$-diversity was observed. The community structure remained even and stable, as evidenced by consistently high J', D, and 1/D indices (**Fig. 3a)**. The microbial composition was consistently dominated by *Propionimicrobium lymphophilum*, followed by a persistent consortium of *Parvimonas micra, Peptococcus niger,* and the *S. anginosus* group, which co-occurred at near-equal

relative abundances. Additional taxa, including *Gemella, Aerococcus,* and *Actinomyces* were also consistently detected as part of this stable, polymicrobial community (**Fig. 3b**).

104 demonstrated higher baseline diversity compared to the three previously discussed dynamic participants. It is also notable that 15 species were detected exclusively in 104 but absent from all dynamic communities, of which 11 were oral anaerobes (**Table 2**). In contrast, 22 species were present in one or more of the dynamic participants but absent from participant 104, including 7 uropathogens (**Table 3**). While participant 204 exhibited high baseline oral microbiome abundance (67.8%), this was dominated by a single species (*Streptococcus anginosus* group), in contrast to the diverse oral consortium in participant 104.

<u>*Participant 206.*</u> This 44-year-old female with a suprapubic catheter had a stable and rich microbial community throughout the study period. Observed richness remained consistent, ranging from 8 to 12 species across 22 weeks. Similarly, H′ indices indicated a persistently diverse ecosystem (H' = 1.25-2.17), with only one transient deviation at week 5 (1.25). This stability was further supported by consistently high D (0.51-0.87) and J′ (0.52-0.90) indices, reflecting a well-balanced community structure (**Fig. 3c)**. Catheter changes occurred at weeks 8 and 19 yet produced no observable disruption to this stable ecological state. The transient dip in all diversity metrics at week 5 occurred independently of any recorded catheter change or antibiotic intervention.

The microbial community in this participant was characterized by co-dominance of metabolically synergistic organisms across the study period [36]. The community was consistently dominated by *Veillonella parvula*, a classic oral commensal and lactatefermenter, alongside known uropathogens *M. morganii*, *P. stuartii*, and *P. aeruginosa*. This may suggest a potential cross-feeding relationship where fermentative byproducts, such as lactate from *V. parvula*, fuel the growth of others persistent species by generating secondary metabolites (organic acids, CO2, H2) [37]. This aligns with established principles of complex microbial communities, where cross-feeding and signaling increase the growth and biofilm formation of one or more species, possibly promoting positive co-occurrences [10, 38]. *P. stuartii* and *M. morganii* alter community virulence and fitness while *P. aeruginosa* is primed for co-colonization in CAUTI settings [39-42]. Other species included the oral cavity-associated *Fusobacterium nucleatum* and *Neisseria elongata*, the ubiquitous nosocomial agent *E. faecalis*, and the skin commensal *Corynebacterium striatum*. The low but consistent presence of *P. mirabilis,* which is known for its role in dense metabolite-structured networks in urine, further supports a stable, synergistic polymicrobial community rather than a random occurrence of independent colonizers (**Fig. 3d)** [3, 39].

*Participant 207.* The longitudinal urine from this 57-year-old female with a Foley catheter revealed a microbial community characterized by stable richness. Observed richness remained consistent ranging from 6 to 10 over 17 weeks, indicating a persistent core set of species. However, the Shannon (H' = 0.66-1.59) and Simpson (D = 0.31-0.76) indices showed fluctuation, pointing to variations in community evenness rather than species loss (**Fig. 3e)**.

This community was less diverse than the others, but it was resilient to catheter changes at weeks 3 and 14, which produced no visible disruption in richness to the community structure. However, notable ecological shifts were observed with a drop in all diversity and evenness metrics at week 4 and a sustained decline after week 14. The week 4 dip was transient, and the community quickly rebounded, suggesting a minor, self-resolving perturbation similar to what was observed in participant 206 at week 5. In contrast, the decline after week 14 that progressed through week 17 was characterized by a collapse in evenness (J' falling from 0.73 to 0.45) while richness held steady.

The community was consistently dominated by a core consortium of *S. aureus, E. faecalis*, and *Actinomyces hominis* present in high, stable proportions. This dominant group was accompanied by a secondary, persistent cluster comprising the oral commensal *Fusobacterium nucleatum* and *Lactobacillus rhamnosus*. Notably, this was the only female participant who harbored a probiotic *Lactobacillus* strain in their catheterized urine specimens. A low-abundance background of *Actinotignum timonense* and *Actinotignum schaalii* was also present in all urine samples (**Fig. 3f)**.

<u>Participant 101</u>. This 77-year-old male with a suprapubic catheter exhibited a diverse baseline community (H′ = 0.83). Catheter changes before study visits 2, 8, 15, 20, 24, and 30 and did not result in significant shifts in $\alpha$-diversity (Kruskal-Wallis test, p = 0.7342). In contrast, the prophylactic administration of Bactrim (Trimethoprim-sulfonamide combination drug) between weeks 8 and 9 due to an accidental catheter removal and reinsertion event with complications led to a sharp decrease in diversity. The H’ decreased from 1.18 at week 8 to 0 at week 9, accompanied by a drop in richness from 8 species to a single taxon. However, this suppression was short-lived; diversity metrics began to recover immediately after antibiotic cessation (H′ = 0.07 at week 10) and returned to pre-antibiotic levels by week 18 (H′ = 1.38) (**Fig. 4a).**

The pre-antibiotic community was initially dominated by *A. urinae (weeks 0-4)*, and then temporarily displaced prior to antibiotic exposure by a consortium of *M. morganii*, *E. coli*, and *P. stuartii* (week 5-8). Bactrim effectively eliminated TMP-SMX-sensitive bacteria but selected for a resistant organism (*P. aeruginosa*) (**Table 4**), which was then displaced by *Providencia rettgeri* (weeks 10-17) [36]. Despite these transient shifts, the community ultimately re-established a new and stable polymicrobial state, dominated by *Propionimicrobium lymphophilum* and *P. stuartii* (weeks 18-30) that was distinct from the baseline *A. urinae* community (**Fig. 4b).**

*Participant 203.* In this 68-year-old male with a suprapubic catheter, a low number of species was detected over the study period. No catheter changes were noted but the participant received oral Keflex (cephalexin) for a skin condition prior to week 7, 8, 9, and 21. S (~2) and H' (0.052) were both low at baseline (weeks 0-5) and remained low throughout the study period, but with fluctuating blooms after cessation of antibiotics (**Fig. 4c**).

At baseline, *P. aeruginosa* (28%) and *P. mirabilis* (72.2%) were detected; *P. mirabilis* was eliminated by oral antibiotic administration between weeks 6-9 while *P. aeruginosa* remained consistent and dominant, likely because *P. aeruginosa* isolates typically exhibit high level resistance to cephalosporins [37, 38]. However, post-antibiotic treatment (weeks 10-20), there was new acquisition of *E. faecalis, Cutibacterium acnes,* and *S. aureus.* Repeated treatment at week 21 exhibited the same pattern with elimination of everything except *P. aeruginosa* (weeks 21-22), and post antibiotic treatment there was emergence of *K. pneumoniae* (30.4%), *K. variicola* (8.6%) and *K. quasipneumoniae* (2.3%) (**Fig. 4d, 4e**).

*Participant 201.* This 59-year-old male with a suprapubic catheter was the only study participant to experience clinician-diagnosed and treated CAUTI, making their urine community dynamics particularly informative. The study period was marked by intense clinical interventions with catheter changes at after weeks 1, 6, 7, 9, 10, 12, and 17, along with 3 distinct courses of antibiotic treatment following CAUTI diagnosis after weeks 3 and 17 (**Fig. 5a).** This participant was also of particular interest for metagenomic analysis as all culture-positive species detected in the urine sample taken immediately prior to each CAUTI episode had been present asymptomatically for at least four weeks [5]. Furthermore, the CAUTI that occurred in week 17 was only one day after the study visit, it was a severe infection that included pyelonephritis (kidney infection) with sepsis, and the same bacterial species cultured from the study visit were also present in the hospital urine and blood cultures [5].

Urine specimens from the baseline visit and first three weeks exhibited moderate diversity (H' = 0.72-1.57). The first antibiotic course of oral ciprofloxacin at week 3 induced a significant but transient collapse, as evidenced by a sharp drop in richness from S=7 to S=3 as well as diversity (H'=1.49 to H'=0.56). This initial perturbation saw the displacement of a core consortium comprising *P. stuartii***,** *Peptoniphilus harei***,** and the *S. anginosus* group (PPS) by a transient bloom of *Staphylococcus aureus* and *Corynebacterium striatum* (**Fig. 5b).**

The community then recovered, with diversity indices and the core PPS consortium rebounding by week 7. This period also saw the new acquisition of *M. morganii*, forming a stable, four-taxa pathogen cluster with a transient increase in *P. stuartii* at week 15 that resolved by week 17. However, a second infection episode occurred one day after the week 17 study visit (**Fig. S3**). Pre-hospitalization, the community was co-dominated by *P. stuartii* (45.1%) and *M. morganii* (10.7%), which together accounted for 55.8% of the total abundance. Post-treatment with Vancomycin/Zosyn followed by Piperacillin/Tazobactam*,* a clear ecological succession was observed where the combined abundance of these two pathogens fell 21-fold to 2.6%, while *S. anginosus* became the dominant successor, increasing to 54.6% relative abundance. All diversity metrics plummeted to zero by week 19 (H'=0, S=0) (**Fig. 5a)***.* Interestingly, all species detected by MetaPhlAn4 at week 17 had been present at similar relative abundances since week 7, indicating that infection onset was not driven by acquisition of a new microbe or a major shift in the resident community. Unlike most participants, whose antibiotic exposures were for non-urinary infections, antibiotic use in participants 201 and 101 was specifically directed at urinary tract management, making their trajectories particularly informative for understanding the ecological consequences of UTI-directed treatment on catheter-associated urinary microbiota.

There was one notable discrepancy between the metagenomic output and our previous culture-based examination of the urine specimens from Participant 201: we previously cultured and isolated a Gram-positive species from the week 22 urine specimen [5], yet MetaPhlAn4 classified this urine specimen as lacking significant microbial reads. A limitation of the MetaPhlAn4 pipeline is that reads are only mapped to curated clade-specific markers. Other tools, such as Kraken2, can instead use k-mers against a broad genome database (RefSeq/PlusPF) to bypass this issue, but can also result in false positives. To compare taxonomic classification methods and determine if potential low-abundance microbes or ones not present in the MetaPhlAn4 database could have been missed by our approach, a comparative analysis of Kraken2 v1.2.1 and MetaPhlAn4 outputs were conducted for participant 201.

After applying a stringent relative abundance threshold (>0.5%), Kraken2 profiles revealed 23 unique microbial species with 6 species at week 22 versus a total of 16 species by MetaPhlAn4. The 0.5% threshold was chosen to focus on taxa with biologically meaningful abundance and to mitigate k-mer artifacts. Importantly, using the 0.5% threshold filtering step reduced the species list from an initial range of 6–362 taxa per sample, with the final 23 species largely representing clinically-relevant uropathogens. Validation of this threshold was supported by two observations: 1) analysis of the engineered positive control sample had false positives when higher thresholds were used, as Kraken2 detected 89 species absent in the engineered community, and 2) 492 of the 543 total species identified across the longitudinal dataset never exceeded 0.1% relative abundance in any sample and were thus classified as artifacts rather than true community members. The 0.5% threshold took care of the false positive outputs and matched our expected species count for the engineered community.

The comparative analysis between the two tools revealed concordance in overall diversity metrics (**Fig. 5c vs 5a**) and community profiles (**Fig. 5d vs 5b**) but identified key differences in taxonomic resolution. Diversity indices and species relative abundances derived from the two pipelines showed strong positive correlations across all samples (r = 0.76-0.92) (**Table S6, Fig. S4**). Kraken2 consistently reported a slightly higher median species richness (~2 species per sample), attributable to its finer taxonomic resolution. The main divergence was in classification. MetaPhlAn4 classified all reads of *Peptoniphilus* as *P. harei*, whereas Kraken2 resolved them into *P. harei* (21%

prevalence) and *P. sp. SAHP1* (68% prevalence). When all *Peptoniphilus* reads in Kraken2 were aggregated, correlation with MetaPhlAn4 was near-perfect (r = 0.98). Similarly, MetaPhlAn4 reported the *Streptococcus anginosus* group as a single clade, while Kraken2 provided species-level identification (e.g., *S. constellatus*, *S. intermedius*) (**Fig. 5b, 5d**).

Collectively, cross-validation between MetaPhlAn4 and Kraken2 confirmed that expanding taxonomic resolution did not reveal additional candidate drivers of infection onset beyond those identified through MetaPhlAn4-based profiling. While Kraken2 provided finer species-level distinctions within select genera, the overall microbial landscape remained consistent across both platforms. Whether deeper species-level resolution within these genera contributes meaningfully to predicting infection sign and symptom onset remains an open question, one that the species associations and clinical outcome analyses presented below begin to address.
